# A model of the appearance of the moving human eye

**DOI:** 10.1101/2021.02.02.429411

**Authors:** Geoffrey K Aguirre

**Affiliations:** Department of Neurology, Perelman School of Medicine, University of Pennsylvania, Philadelphia, PA 19104

## Abstract

The entrance pupil and first Purkinje reflection (“glint”) in an image of the human eye serve as important features in 3D, model-based eye tracking applications. Here I present a physically and biologically accurate, ray-traced model that supplies the appearance of the pupil and glint of a moving eye. Once biometrically calibrated for a subject under study, simultaneous fitting of the pupil and glint features by the model supplies eye rotation, radius of the aperture stop, and the translation of the eye relative to the initial camera position. The biometric parameters of the model, including corneal curvature and the depth of the centers of rotation, are obtained by model-based fitting of images of the eye posed at known gaze angles. This approach was applied to eye recordings from 30 people obtained during gaze calibration, and while head position was recorded simultaneously using echoplanar magnetic resonance imaging. The refractive and reflective effects of spectacle and contact lenses worn by some subjects were incorporated into the model. The fitted parameters reveal that the center of rotation for horizontal eye movements is deeper (13.8 mm) than that for vertical eye movements (12.1) mm, consistent with prior studies. Individual differences in the depth of the rotation centers, and in corneal curvature, were well related to biometric measures obtained from these subjects using clinical ophthalmologic instruments. Once biometrically calibrated for each subject, gaze position was modeled with a cross-validated, median (across subject) absolute error of 0.58°, and image-plane translation of the head was estimated with sub-millimeter accuracy. The open-source model described here produces biologically accurate simulations of the appearance of the eye in motion, and may be used in model-based search to derive eye pose and biometric properties from empirical data.

## Introduction

The measurement of eye movement and pupil response is central to investigations in neuroscience and psychology, and in the engineering of devices with a human-machine interface. There is a rich history of methodological approaches to measuring eye movements, with non-contact, optical techniques preeminent. Most commonly, one or more cameras—imaging in the visible or infrared—are used to observe the eye of a subject, often with the eye illuminated with one or more light sources. A central analytic challenge is then to derive the pupil size and gaze position (also termed the point-of-regard) of the subject from these images. While numerous approaches have been advanced over the decades, feature-based, 3D model techniques have seen the greatest development. In this class of analysis approach, features from the image are extracted, and then the shape and position of the features are fit by reference to a simplified model of the eye that might have produced the image. One such feature is the entrance pupil of the eye, which is the virtual image of the aperture stop of the iris as seen through the refractive effects of the cornea. A second important feature is the position of one or more “glints”, which are the reflection of a light source by the tear film of the cornea (also known as the first Purkinje image). Gaze position and pupil size are then derived via a system of equations that approximates the geometric properties and refractive effects of a rotated eye upon these features.

While computationally expedient, these approaches often have limited generalizability. In most cases, analytic solutions attempt to measure one property of the eye under study (e.g., gaze position) while discounting the effect of other properties that are not explicitly estimated (e.g., pupil size or head translation). When implemented with simple imaging systems, analytic approaches become fragile. For example, small head movement or inaccuracy in the modeling of biometric variation can substantially limit single-camera imaging systems (Guestrin 2006). Additional, ad-hoc corrections to analytic approaches are required to correct for the many imperfections and complications that can accompany an eye recording (e.g., corrective lenses worn by the subject, or partial occlusion of an eye feature by the eyelid; Hansen 2009).

An alternative to analytic approaches is to create a biologically accurate, “forward model” of the eye under study. Guided by parameter settings, a forward model may be used to generate simulations of the appearance of eye features. The output of the model will naturally capture complicated, non-linear interactions of the parameters that are difficult to reduce to an analytic form. Model output can be compared to empirical observations, with a search across parameter values used to recover the properties and position of the eye under study. Beyond improving the accuracy of model-based eye tracking approaches, a biologically accurate, forward model of the appearance of the eye can be used to provide synthetic input to the training of appearance-based eye tracking systems (Swirski 2014; Yiu 2019), and support measurement of biometric properties of the eye.

Here I develop a forward model of the appearance of the moving human eye. A ray-traced model of the entrance pupil of the eye (Aguirre 2019) is supplied with biologically accurate rotation, and a solution for the appearance of one or more glints produced by an active imaging light source. Parameters of the model control relevant sources of biometric variation, including the shape of the front surface of the cornea, and the centers of rotation of the eye. To validate the model, I show that the forward output may be harnessed to support model-based fitting of image features from an eye under observation. While applicable to almost any combination of imaging equipment, I focus here upon the commonly encountered circumstance of observing an eye using one camera and one light source (OCOL). I evaluate the ability of model-based search under these conditions to recover accurate biometric values, gaze position, and head translation for an observed eye. In simulations, I demonstrate that these parameters are well estimated, as long as the distance of the camera from the eye is known. I then analyze empirical data collected from 30 subjects using an OCOL imaging system. When supplied with observations of the eye posed at known angles, the model recovers biometric parameters that are well matched to measures from clinical ophthalmologic instruments. Eye recordings were also made from these subjects during simultaneous collection of echoplanar functional MRI data at 800 msec resolution, providing an independent measure of head translation. When supplied with biometric parameters, the model recovers gaze position with a median absolute error of 0.58°, and recovers in-plane translation of the head with sub-millimeter accuracy.

### Model eye description

The approach begins with a previously described, forward-model of the appearance of the human eye implemented in software (Aguirre 2019). The software describes the eye as a set of quadric surfaces (ellipsoids and hyperboloids), each with an index of refraction that is set for the wavelength domain used to image the eye (e.g., visible or infrared). The position and intrinsic properties of a camera are also defined, and ray tracing is used to obtain the image of eye features as seen by the camera (Figure 1a).

**Figure 1.**
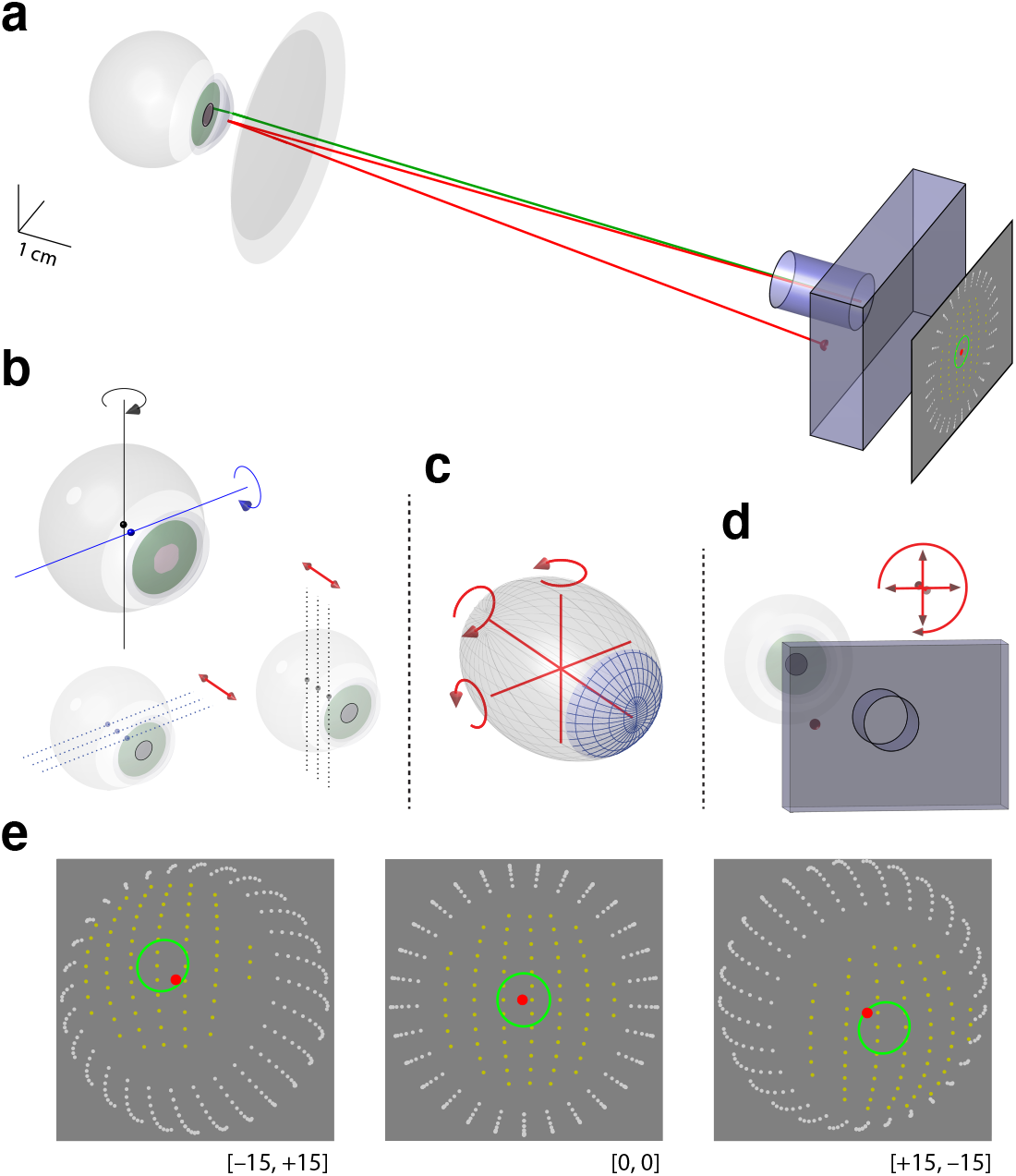
Forward model of the human eye in rotation. **a)** The eye is described by a set of refractive quadric surfaces. The appearance of the eye in the camera image is obtained by ray tracing points from the eye to the nodal point of the camera. Points from the border of the aperture stop of the iris (green) are subject to the refractive effects of the cornea. The first Purkinje image is the point of reflection of an active light source (red) off the tear film of the eye that intersects the camera nodal point. The model may include artificial lenses worn by the subject, such as the –5 diopter ophthalmic spectacle lens shown here. **b)** Parameters of the model control the depth of the center of rotation of the eye for horizontal (black) or vertical (blue) movements. **c)** The front surface of the cornea (blue) is modeled as an ellipsoid (gray). Six parameters control the semi-axis lengths and the rotation of the ellipsoid. **d)** Four parameters control the translation and torsion of the camera with respect to the eye. **e)** Given biometric and camera position parameters, the model provides the appearance of the eye features in the camera image for different poses of the eye. Shown are three poses of the eye with varying [horizontal, vertical] rotation, all with an aperture stop radius of 2mm. The border of the entrance pupil is described by an ellipse (green line). The locations of any glint(s) are indicated as well (red point). The front surface of the cornea (yellow) and the ellipsoidal vitreous chamber (white) are also shown for purposes of illustration.

One of these features is the pupil. Rays that arise from the edge of the aperture stop of the iris are traced through the cornea to the nodal point of the camera (green line). A set of these points are used to define the form of the pupil ellipse in the camera image. The ray-trace solution can include corrective lenses. These are defined by their spherical optical power, and then added to the model at either an appropriate vertex distance (for a spectacle lens), or shaped to match the curvature of the modeled cornea (for a contact lens); see the Supplementary Materials for more details.

A second feature is the “glint”, or first Purkinje reflection. The position of active light sources are specified relative to the nodal point of the camera. A glint is identified by a ray that departs the light source, is reflected by the tear film of the corneal surface, and returns to the camera (red line). When a contact lens is included in the model, the glint is returned from the tear film on the contact lens.

The appearance of the pupil and glint features in the simulated camera image is influenced by the biometric parameters used to model the eye, the position of the camera, and the “pose” of the eye. The pose specifies rotation of the eye with respect to the camera, and the radius of the aperture stop. Table 1 summarizes the parameters used to describe the appearance of the pupil and glint in a moving eye. The entire forward model (Aguirre 2019) includes many more parameters that define the eye (e.g., vitreous chamber size, properties of the crystalline lens, corneal thickness), but which have minimal influence upon the appearance of the eye features we consider here.

**Table 1.**
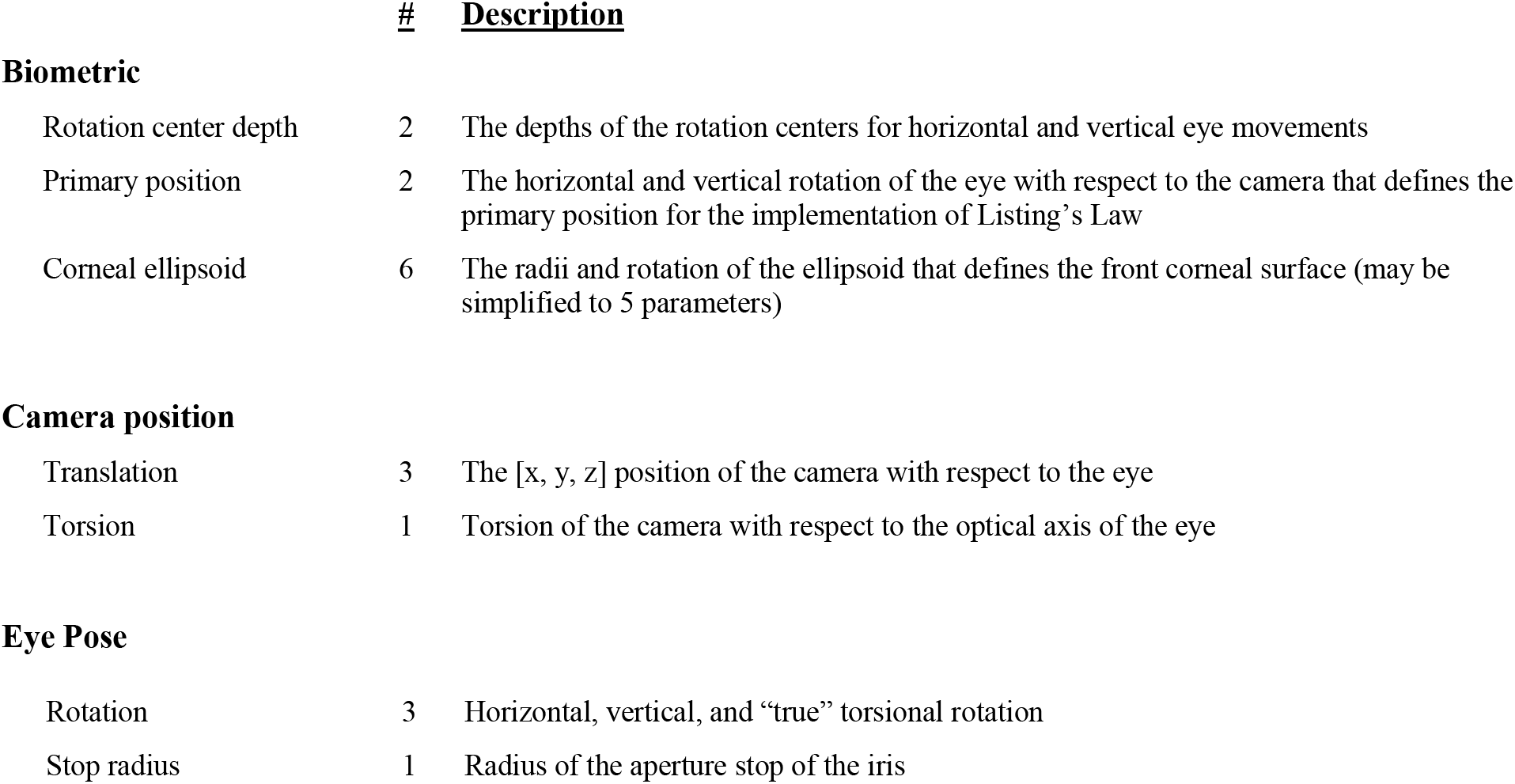
Parameters of the model. In the current work we will focus upon 18 parameters that influence the appearance of the pupil and glint, arising from a description of eye biometry, camera position, and the dynamic “pose” of the eye.

|  | # | Description |
| --- | --- | --- |
| **Biometric** |  |  |
| Rotation center depth | 2 | The depths of the rotation centers for horizontal and vertical eye movements |
| Primary position | 2 | The horizontal and vertical rotation of the eye with respect to the camera that defines the primary position for the implementation of Listing’s Law |
| Corneal ellipsoid | 6 | The radii and rotation of the ellipsoid that defines the front corneal surface (may be simplified to 5 parameters) |
| **Camera position** |  |  |
| Translation | 3 | The [x, y, z] position of the camera with respect to the eye |
| Torsion | 1 | Torsion of the camera with respect to the optical axis of the eye |
| Eye Pose |  |  |
| Rotation | 3 | Horizontal, vertical, and “true” torsional rotation |
| Stop radius | 1 | Radius of the aperture stop of the iris |

#### Biometric properties

While the rotation center of the eye is often treated as a single point (von Noorden 2001), there are in fact separate centers for vertical and horizontal rotation (Fry & Hill 1962, 1963). Fry & Hill found that the depth of the center of rotation (relative to the corneal apex) was on average greater for horizontal eye movements as compared to vertical eye movements. Further, the horizontal center of rotation was found to be slightly nasal to the optical axis (Figure 1b, left). The precise depth of the center of rotation is found to vary across individuals, with this variation correlated with the axial length of the eye (Dick 1990). Therefore, model parameters allow the depth of each rotation center to be scaled separately for the eye under study (Figure 1b, right).

Eye movements (in the absence of head rotation) generally obey Listing’s Law, which observes that there is no change in the “true” torsion of the eye (with respect to its optical axis) with a change in position. When 3D rotations are implemented with quaternions, this property is automatically achieved. In the current model, however, I implement eye movement as a series of rotations following “Fick coordinates” (Haslwanter 1995), as this allows specification of separate centers for each direction of eye rotation. To implement Listing’s Law, the model adds “pseudo-torsion” about the optical axis of the eye (Nakayama 1983). The calculation of pseudo torsion is performed relative to the “primary position” of the eye, which is the position from which any other eye pose could be achieved by rotation of the eye about a single axis (if the horizontal and vertical rotation centers were identical). The primary position is defined by two parameters (horizontal and vertical eye rotation with respect to the camera) which may be set for the eye under study.

The shape of the front corneal surface influences both the appearance of the entrance pupil and the imaged position of the glint (Figure 1c). The model describes the front corneal surface as a tri-axial ellipsoid, defined by the lengths of the three semi-axes, and the rotation of the ellipsoid (torsion, “tilt”, and “tip”) relative to the optical axis of the model eye. The initial parameter values for the ellipsoid are taken from population average measurements (Navarro 2006), but can be adjusted by calibration for the eye under study. As the depth of the anterior chamber is set by a separate parameter, it is possible to find different combinations of semi-axis lengths that produce models of the corneal front surface with nearly identical curvature. To avoid this over-parameterization, the axial length of the ellipsoid may be held constant at a default value, and the curvature of the corneal front surface expressed in the familiar values of keratometric diopters, k1 and k2. The k1 and k2 values describe optical power at the flattest and steepest meridians (respectively) of the front surface of the cornea. These values are in units of diopters, expressed in terms of a particular, standard refractive index for the anterior surface of the cornea that incorporates the refractive effect of the back corneal surface as well. Specified values of k1 and k2 are converted within the software into corresponding ellipsoidal semi-axis lengths to be realized within the model.

#### Camera position

The position of the camera (Figure 1d) is defined by its translation and torsion relative to the corneal apex when the optical axis of the eye and the optical axis of the camera are aligned (and when the corneal ellipsoid is also aligned with the optical axis). Because eye rotation is defined relative to this frame, specification of the pitch and yaw of the camera is not necessary. As a matter of convenience, we can consider camera position both as an initial, absolute position, and also as relative change in that position from moment to moment. In the model coordinate system, a translation of the head or eye is equivalent to a change in relative camera position.

#### Eye pose

Given parameters that define the eye and camera position, the appearance of the pupil and glint may then be obtained for a given eye pose. The eye pose is defined by the horizontal, vertical, and true torsional rotation of the eye, and the radius of the aperture stop of the iris. Figure 1e presents images of the appearance of eye features (for the eye and camera shown in Figure 1a) when the eye is posed with different combinations of horizontal and vertical rotation, with true torsion and the stop radius held constant.

#### Deriving pose and translation from image features

One application of the model is to support a search to estimate the values of an unknown eye pose and translation for an observed pupil and glint. Consider the case in which we already know (through some means) the biometric parameters of an eye under study. Given an image of the eye (Figure 2, left), we can use any of a number of segmentation approaches (Fuhl 2016) to identify the location of the glint and points that are on the perimeter of the entrance pupil (Figure 2, center). We might then search across values of camera position and eye pose to find a modeled pupil perimeter and glint location that best matches the observed eye features (Figure 2, right).

**Figure 2.**
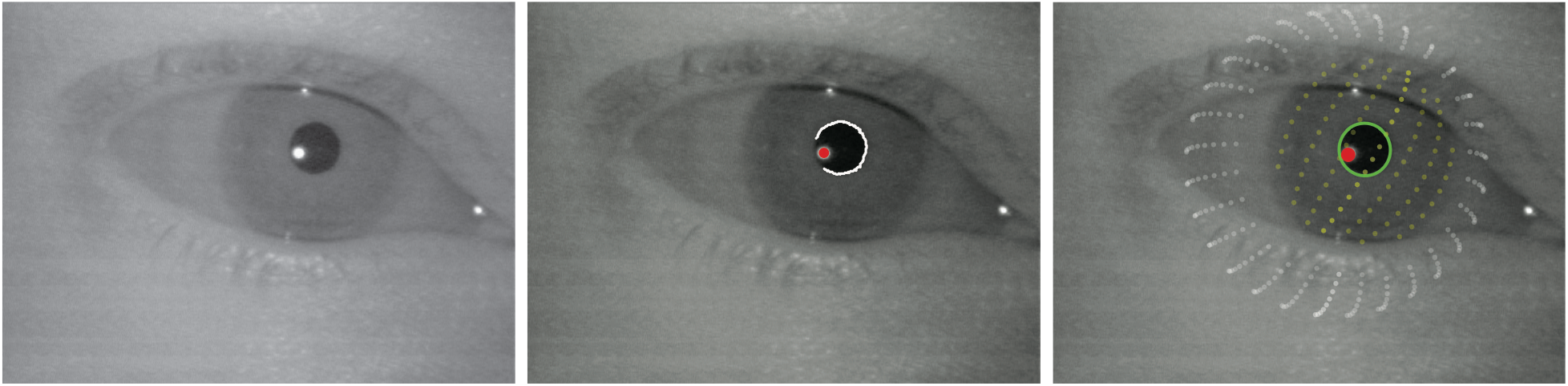
Model-based search to recover eye pose. Given an image of the eye (left), any one of several methods might be used to identify points on the border of the pupil (white) and the location of the glint (red) (center). Given biometric parameters, model-based search may be used to fit these eye features and recover an estimate of the pose of the eye and the relative translation of the camera (right). The modeled location of the glint (red) and the elliptical form of the entrance pupil (green) are shown. The cornea front surface (yellow) and ellipsoidal vitreous chamber (white) are also shown for the purpose of illustration.

In the simulations and empirical tests that I examine here, the search is not conducted over true torsion of the eye; instead this parameter is locked and held constant. This simplification is motivated by practical and theoretical properties of the system under study. Eye torsion has only a small influence upon the appearance of the pupil perimeter. Further, for the head-fixed eye movements studied here, Listing’s Law holds that true eye torsion will not vary with eye position. For these reasons as well, I do not explore the effect of modeling variation in the primary position of the eye.

Given observed points on the pupil perimeter, the software implements a bounded, non-linear search over the remaining parameters (camera translation, horizontal and vertical eye rotation, and the stop radius) with the objective of minimizing the L2 norm between the pupil perimeter points and the simulated pupil ellipse provided by the model (the green ellipse and the white points in Figure 2). This search is further subjected to a non-linear constraint that the simulated and observed location of any glints must match (within a tolerance of less than a pixel). Supplementary Movie 1 illustrates a model-based search of eye features to derive pose and camera position.

#### Relating eye pose to gaze position

In many cases, we wish to derive from an observation of the eye the point in the world where the eye is currently fixating. To do so, we require a measurement of the fixation pose of the eye, and the torsion of the world coordinates with respect to the horizontal plane of rotation of the eye. These parameters may be obtained by observing the eye during a simple calibration in which the subject views a target at known world coordinates. The fixation pose is defined as the rotation angles of the eye (relative to the camera) observed when the gaze of the subject corresponds to a fixation point at the world origin location (i.e., fixated upon a target at zero degrees visual angle). If the camera is optically positioned at this origin point, with zero torsion between the eye, camera, and world coordinates, then these fixation angles are equivalent to the horizontal and vertical angle between the line of sight and optical axis of the eye (Atchison & Smith 2000).

If the subject is wearing a spectacle lens, then the visual world is subjected to magnification/minification, which scales the mapping of eye rotation to degrees of visual angle of the stimulus. This spectacle magnification factor is incorporated into the forward simulation, and is obtained by ray-tracing through the model eye optical system with and without the artificial lens (Figure S1). While contact lenses also introduce a (smaller) magnification factor, a contact lens rotates with the eye, so the angle of rotation and the visual angle of the target remain matched.

In these analyses I treat the rotation angle of an eye between two fixation targets as equivalent to the difference in visual angle between those targets. This is not strictly accurate, as the optical center of the eye (the approximation to the nodal points; Harris 2010) is anterior to the centers of rotation (Steinman 1982). This effect of ocular parallax, however, is small (<0.1 degree) for the case considered here in which the fixation targets are over a meter distant from the eye (Bingham 1993).

### Simulations

We may validate the model-based search approach by attempting to recover the parameters of an eye used to generate simulated eye features. In simulation we first examine the ability of model-based search to recover the rotation and translation of an eye for which the biometric parameters are known, and then the ability to recover biometric parameters when the gaze position is known.

#### Simulated recovery of eye pose and camera translation given biometric parameters

We first consider the situation in which the biometric parameters of an eye under study are known and we wish to recover the pose of the eye and the position of the camera. I examined three simulation cases to characterize this behavior. In all three cases, a model eye with the default biometric parameters is assumed, with this eye observed by a camera with a specified intrinsic matrix (sensor resolution 640 x 480 pixels) and a known initial position. Each case includes 1000 evaluations of the ability of the model to recover a set of randomly selected, simulated eye poses and relative changes in camera position. The relative changes in camera position might occur, for example, if the subject under study makes head movements during the observation. For the particular set of parameters to be tested, the model was used to simulate the coordinates of the glint(s) and 10 points around the pupil perimeter. A moderate amount of normally distributed noise was added to these coordinates (0.25 pixel SD; roughly 2.5x the noise present in the imaging system itself, Guestrin 2006). The non-linear search routine was then applied to the simulated glint and pupil features, and the agreement between the simulated and recovered eye pose and camera translation parameters was recorded.

The first simulation features a camera equipped with two active IR light sources, positioned 14 mm on either side of the camera lens. Figure 3, left, presents the error in the recovered parameters across all simulations. The median absolute error in parameter recovery in these simulations is reported in Table 2. An exception to the overall good performance of the search is found in the recovery of relative camera depth, for which the median error was many times larger than the error in recovery of in-plane camera translation.

**Figure 3.**
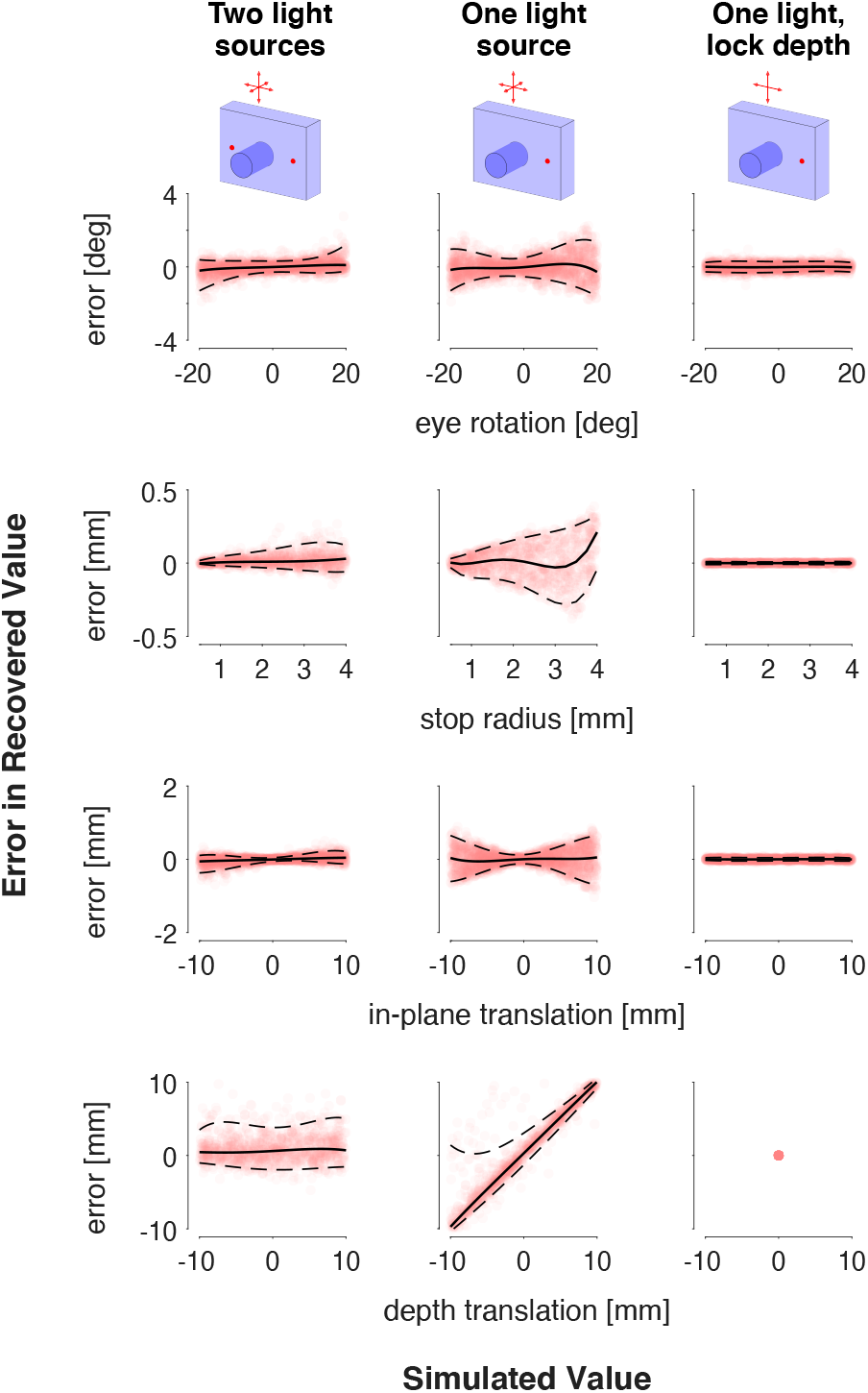
Recovery of eye pose and camera translation in simulation. A default set of biometric parameters was assumed, and eye features (pupil perimeter and glint) were generated for a range of eye poses and camera positions. Each plot provides the error in the recovery of the eye pose or camera position value as a function of that value. One-thousand different sets of pose and camera position values were examined for each of three simulated imaging arrangements (organized by column). The dashed lines provide the 95% confidence interval around the set of1000 values (transparent red points). The plots for eye rotation include data points for both recovered horizontal and vertical rotation. Similarly, the plots for in-plane translation include data points for both horizontal and vertical translation.

**Table 2.** Errors in model-based search. The ability of the model-based search to recover eye pose and relative camera position was examined in simulation. These simulations all featured one camera, with this camera equipped with either two light sources (OCTL) or one (OCOL). A third simulation locked the simulated camera depth and did not search across this parameter.

| Median absolute error | <u>OCTL</u> | <u>OCOL</u> | <u>OCOL (lock depth)</u> |
| --- | --- | --- | --- |
| Eye rotation [deg] | 0.27 | 0.52 | 0.20 |
| Stop radius [mm] | 0.02 | 0.07 | 0.00 |
| In-plane translation [mm] | 0.07 | 0.25 | 0.03 |
| Depth translation [mm] | 1.10 | 4.67 | n/a |

The difficulty in recovering camera depth is exacerbated if the simulation is conducted with a single light source instead of two (Figure 3, middle). Errors are increased in the recovery of all parameter values, but especially so for depth, with the search effectively unable to detect changes in the distance of the camera from the eye. This error in recovering depth translation drives the errors in estimation of the other parameters: the absolute value of the errors in estimation of eye rotation, stop radius, and in-plane translation are all correlated with the simulated change in camera depth (Pearson’s R>0.6). If camera depth is held constant (Figure 3, right), the performance of the search is restored for the OCOL condition. Now, eye rotation, size of the aperture stop, and in-camera-plane translation are estimated with great accuracy.

It has previously been shown that a one camera, two light source configuration is the minimum possible for recovery of both translation and rotation values of an observed eye (when this estimation is made using the pupil center and glint; Shi 2000; Guestrin 2006). The current simulations agree with this observation, and further illustrate the particular challenge of recovery of depth information. If depth is held constant, the minimal system may be simplified to one camera and one light source. Under these conditions, the model-based search implemented here is able to recover the rotation and translation of the eye with precision. In the remaining studies we will examine and validate the model within the constraints of the OCOL configuration.

#### Simulated recovery of biometric parameters given eye pose

We next examine the ability of an OCOL configuration to recover the unknown biometric parameters of an eye under study. This simulation considers that we have observed an eye while it was posed to view nine targets at known visual angles (Figure 4a). This set of nine observations is termed here a “gaze calibration” measurement. Each of the simulated measurements assume different, randomly selected parameters for the in-plane translation of the camera, the rotation centers of the eye, and the curvature of the cornea. In this simulation I did not vary the rotation of the ellipsoid that defines the corneal front surface. I also held constant other aspects of the model related to torsion (primary position of the eye, camera torsion), and camera depth.

**Figure 4.**
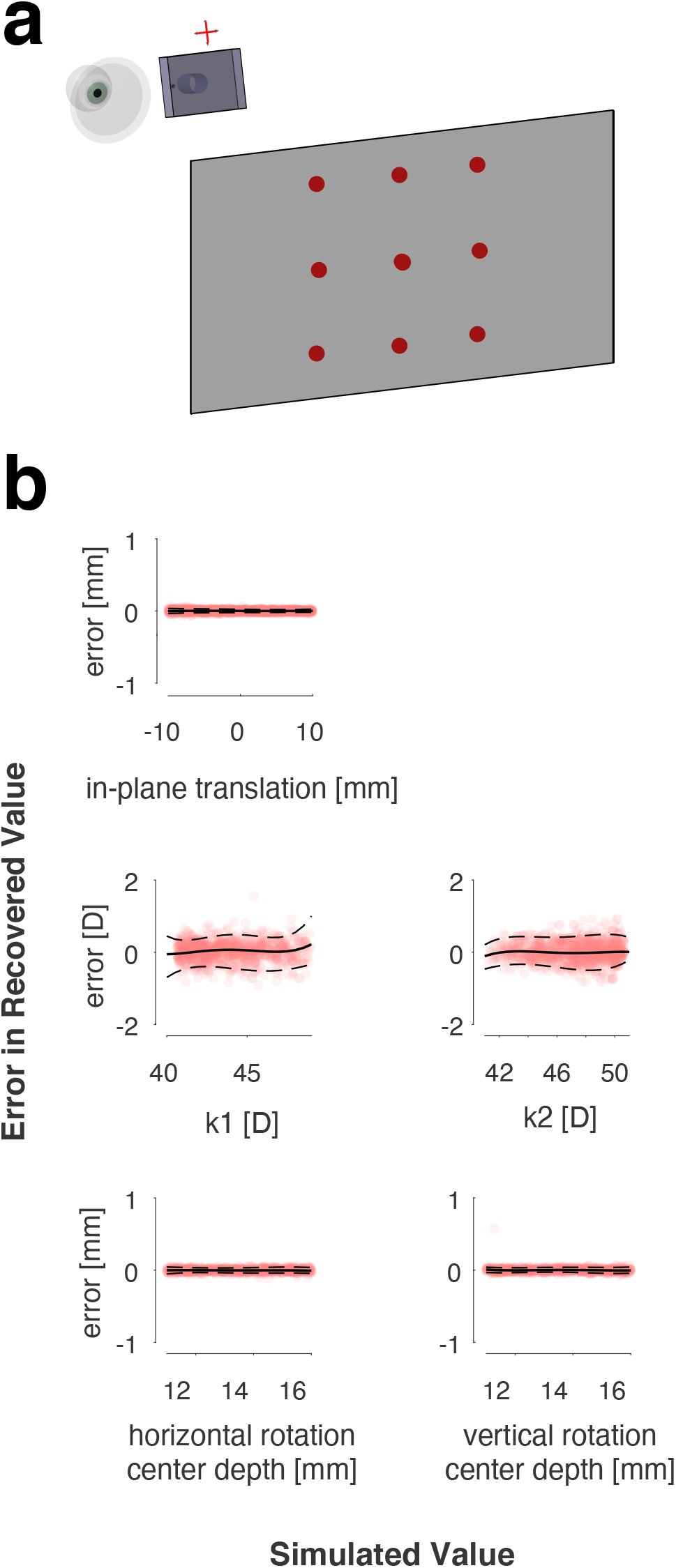
Recovery of biometric parameters in simulation. **a)** In each of one-thousand simulations, a set of biometric values were chosen and the modeled eye was posed to fixate upon nine targets at known visual angles presented on a distant screen (1065 mm away; here illustrated as 355 mm). The pupil and glint features from each of these poses were obtained, and then fit in a model-based search in an attempt to recover the underlying biometric parameters. **b)** Each plot presents the error in recovery of a biometric parameter as a function of the simulated value of that parameter. The dashed lines provide the 95% confidence interval around the set of 1000 values (transparent red points).

### Simulated Value

Each of these simulations provides the perimeter of the pupil and the coordinates of the glint for the nine poses of the eye (with 0.25 SD of normally distributed noise added to these coordinates). These features are then fit in a search across the camera translation and eye biometry. To do so, we search across values of these parameters, and evaluate the ability of the model output to simultaneously fit the perimeter of the pupil, the position of the glint, and the relative rotation of the eye. The search attempts to minimize disagreement between the assigned rotations of the eye and the positions of the gaze targets in degrees of visual angle, while simultaneously minimizing error in fitting the perimeter of the pupil and the location of the glint. Supplementary Movie 2 illustrates a search of this kind.

Figure 4b summarizes the performance of the model-based search across 1000 simulated gaze calibrations. The parameters are recovered with high accuracy. Notably, separate horizontal and vertical rotation centers are found even in the setting of an astigmatic cornea. Across all of these simulations, the estimated camera translation and eye biometry parameters also map eye rotation to gaze position with a median error of 0.03 degrees. These simulations demonstrate that, when camera depth and modeled torsion are held constant, a one camera one light source imaging system can recover eye biometry and camera position from a gaze calibration measurement.

We turn now to the performance of the model in fitting empirical data.

## Experimental Methods

### Subjects and data collection environment

Data were collected from 30 normally sighted participants, recruited from the University of Pennsylvania and the surrounding Philadelphia community. This is the complete subset of a total of 50 subjects (Chen 2020), studied as part of a pre-registered project (https://osf.io/ervrh), who had brain imaging performed (40 subjects) and from whom four gaze-calibration measurements were made during their second imaging session (30 subjects). Participants were required to be at least 18 years of age, have no history of ophthalmologic disease, and have corrected visual acuity of 20/40 or better. There were multiple sessions of data collection for each participant, typically across multiple days. These sessions included retinal imaging, as well as structural and functional magnetic resonance imaging of the brain (MRI). Fourteen of the subjects were women, and the mean age of all participants was 28. This study was approved by the University of Pennsylvania Institutional Review Board, and all participants provided written consent.

Eye recordings were made as part of the MRI study. The head of the subject was positioned in a 64-channel head coil, within a 3 Tesla Prisma scanner (Siemens, Germany). Padding was placed on the sides of the head to encourage the subject to remain motionless, although head motion was still possible. MRI and eye video acquisitions were rejected and recollected at the time of scanning if large or sudden head motion was seen either by video monitoring or from inspection of reconstructed MR images, following the pre-registered protocol. Subjects viewed stimuli presented on an MR-compatible LCD screen (SensaVue fMRI, Invivo corporation) placed at the end of the scanner bore at an optical distance of ~1065 mm from the eye of the subject.

Subjects underwent autorefraction (Canon RK-F1 Full Auto Ref-Keratometer; Canon Medical Systems, Otawara, Tochigi, Japan). The axial length of both eyes was measured in all participants using the IOLMaster 500 (Carl Zeiss Meditec, Jena, Germany). This instrument was also used to obtain keratometric measurements for 24 of the 30 subjects. These data have been presented previously (Chen 2020).

Ten of the subjects wore artificial lenses during data collection to correct for myopia. Seven subjects wore contact lenses, and three subjects wore MR-compatible spectacle lenses. The refractive power of these lenses was assumed to be equivalent to the spherical refractive error of the right eye of the subject as measured by autorefraction, unless a precise value for the optical prescription was available.

### Infrared video recording of the eye, and MR imaging of the brain

An MR-compatible, IR video camera (LiveTrack, Cambridge Research Systems) was placed behind an angled cold mirror, and both the camera and mirror were mounted to the head coil. The mirror both hid the camera and allowed the subject to view the LCD screen. The camera had one active IR LED, located 14 mm lateral to the center of the camera lens. Recordings of the eye were made in NTSC DV 30Hz, and in subsequent processing converted to progressive 60 Hz videos, using “bob” deinterlacing.

The intrinsic camera matrix and radial lens distortion for the camera were measured using the MATLAB camera calibrator application (Bouguet 2012).

Two types of eye recordings were made. During each of four “gaze calibrations,” the subject was asked to fixate upon targets presented on the screen. There were nine targets, arranged in a square, 3×3 grid that was 14° wide. The targets were presented in a random order and the subject was asked to fixate upon each target for several seconds. During each of four “movie watching” acquisitions the subject engaged in free viewing of animated Pixar movie shorts while functional MRI data were acquired. Each of these acquisitions was 336 seconds in duration. The gaze calibration and movie watching acquisitions were interleaved with each other, and with other functional MRI acquisitions not discussed here. The scanning session was 60 minutes in duration. Subjects were permitted to make small head movements between acquisitions. The thirty subjects examined here completed all gaze calibration acquisitions; other subjects were unable to complete the full measurement set due to discomfort, fatigue, or various technical failures of the eye recording system.

Echoplanar functional MRI data were collected during movie watching using the Human Connectome Project (HCP) LifeSpan protocol (VD13D). BOLD fMRI data were obtained over 72 axial slices with 2 mm isotropic voxels with multi-band = 8, TR = 800 ms, TE = 37 ms, FOV = 208 mm, flip angle = 52°. A T1-weighted, 3D, magnetization-prepared rapid gradient-echo (MPRAGE) anatomical image was also acquired for each subject in axial orientation with 0.8 mm isotropic voxels, repetition time (TR) = 2.4 s, echo time (TE) = 2.22 ms, inversion time (TI) = 1000 ms, field of view (FoV) = 256 mm, flip angle = 8°. A TTL pulse produced by the scanner at the start of each acquisition was recorded and used to synchronize the fMRI and eye video data.

### Pre-processing of eye videos and definition of eye features

The focus of the current report is upon the analysis of eye features, and the approach described is agnostic as to the method used to derive those features from observations of the eye. I provide here a brief overview of the pre-processing stage of the eye videos. The full data analysis toolbox is available as open source code (https://github.com/gkaguirrelab/transparentTrack), as is the code that contains the precise parameters used to process the data presented here (https://github.com/gkaguirrelab/eyeTrackTOMEAnalysis).

Following deinterlacing, eye videos were processed to identify the location of the glint and the perimeter of the pupil. The glint location on each frame of the video was found as the centroid of the region with the highest (brightest) image intensity. The pupil perimeter was initially identified in each frame using the Hough transform, as implemented in the *imfindcircles* function in MATLAB. In most cases, this initial segmentation included imperfections, either because the pupil was in shadow, the border between the pupil and iris was indistinct, or the pupil border was obscured by the eyelid, eyelashes, or glint. To correct these imperfections, the perimeter of the pupil underwent iterative cleaning, which included masking out the effect of the glint and cutting away portions of the perimeter. This cleaning attempted to maximize the extent of the retained pupil perimeter while minimizing the error in the fit of an ellipse to the remaining perimeter points.

For the gaze calibration acquisitions, a representative video frame was selected for each fixation target. An automated procedure first selected those frames with error in the fitting of an ellipse to the pupil perimeter. From amongst these, the frame was identified for which the center of the pupil ellipse was closest to the median position of pupil centers across frames for that fixation target. In a small number of cases, this automated routine failed to select an appropriate frame, in which case an alternate frame was selected by hand.

### Derivation of eye pose from eye features

I developed a non-linear search algorithm to derive camera translation and eye pose (rotation angles and size of the aperture stop) from the pupil perimeter and glint location observed in a video image frame. The search begins with a model eye defined in software (including the biometric parameters listed in Table 1) and specification of the properties and initial position of the camera. A forward projection of the model eye (Aguirre 2019) is used to generate a prediction of the appearance in the image plane of the pupil ellipse and glint location for a candidate camera position and eye pose. This prediction includes the elliptical form of the pupil, and the search across parameter values attempts to minimize the L2 error in fitting the pupil perimeter points with the pupil ellipse. The search is also subject to the non-linear constraint that the predicted and observed glint location(s) in the image match to within 1 pixel distance. The matching of observed and predicted glint location is treated as a constraint (as opposed to objective) as I observed in empirical data that there was very little noise in the measurement of glint location. This search, including placing bounds upon the parameters, is conducted using *fmincon* in MATLAB.

### Derivation of biometric parameters from gaze calibration data

A separate non-linear search algorithm was developed to identify the biometric and initial camera position parameters that best account for the set of eye features observed during a gaze calibration. This fitting procedure for gaze calibration data is essentially a wrapper around the eye pose search described above. On each iteration of the gaze calibration search, the set of eye features is fit (using the eye pose search described above) using candidate values for initial camera position and eye biometry (rotation centers, corneal curvature, corneal rotation, eye primary position). The fit to the set of eye features yields four different error metrics:

1. The L2 norm of the errors in fitting the pupil perimeter across the fixations.
2. The L2 norm of the errors in fitting the glint location(s) across the fixations (these values generally remain close to zero given that the eye pose search imposes this as a constraint).
3. A camera displacement error. The fit to each fixation yields a measure of change in camera position from the candidate initial position. If the candidate camera position is correct, and the eye features from each fixation reflect a pure rotation of the eye without translation, then the change in relative camera position for each fixation should be zero. Small displacements of the eye with rotation may be present due to the kinematics of eye rotation, or from small head movements; large displacement values may reflect a poorly fitting biometric model. The camera displacement error metric is the L2 norm of the magnitude of relative camera translations across fixations, scaled by the direction of these translations. Thus, this error metric grows when the eye pose solutions include large camera translations, and further grows when those translations are in a consistent direction. Reduction in this error may therefore be achieved by finding an initial camera position that is centered amongst the positions of the set of eye features.
4. A gaze error. The set of eye poses may be mapped to the set of fixation targets by a three-parameter affine transformation (corresponding to the fixation pose of the eye, and the torsion of the screen relative to the horizontal plane of rotation of the eye). Having done so, we obtain the L2 norm of the disagreement between the set of eye pose angles and the visual angles of the fixation targets (adjusted for spectacle magnification where necessary), yielding a gaze error metric.

A weighted combination of these four error metrics yields an omnibus error for the candidate biometric and camera position parameters in fitting the gaze calibration eye features. I conduct a bounded, non-linear search across these parameters using Bayesian Adaptive Direct Search (Acerbi 2017). To avoid local minima, the search is conducted in stages, with different sets of parameters undergoing optimization at different stages. Generally, those parameters with the largest effect upon the fitting of the features (e.g., eye rotation depth) undergo optimization earlier, and those with the smallest effect (e.g., primary position of the eye) undergo optimization later.

Two parameters of the search—camera distance and camera torsion—were set to initial values by hand for each gaze calibration acquisition. Although the camera was mounted at a fixed point on the head coil, individual variation in head size produced variation in the distance of the camera from the eye of the subject. An initial value for camera depth was obtained by adjusting this parameter and examining the appearance of the model eye superimposed on a video frame for the subject, with the goal to match the modeled and observed sizes of the iris, vitreous chamber, and inter-canthal distance. Camera torsion was also estimated by hand. In most people there is a positive canthal angle, meaning that the medial canthus is slightly lower on the face than the lateral canthus. The principle source of biometric variation in this angle is the racial appearance of the face (Rhee 2012). In the current study, a measurement of the canthal angle, referenced to the race of the subject, was used to derived an initial value for camera torsion. These parameters were allowed to vary in the gaze calibration search, although changes from their initial values contributed to the omnibus error metric. This regularization was necessary as the model has little purchase on these parameter values in the setting of a one camera, one light source imaging arrangement.

Multiple gaze calibration acquisitions from a subject were fit simultaneously. To do so, the biometric parameters were held in common across acquisitions, while the camera position parameters were free to vary. The omnibus errors from each gaze calibration acquisition were combined into a multi-acquisition objective using an L1 norm.

The resulting biometric parameters and camera position information was used to derive eye pose values from the movie watching acquisitions for each subject. Additionally, the biometric parameters derived from the gaze calibration data were compared to the biometric measurements made with ophthalmologic instruments on those same subjects.

### Derivation of relative camera position from brain motion

The MRI data were processed using the HCP minimal preprocessing pipeline (Glasser 2013). A component of this processing is correction of the functional data for head motion (FLIRT; Jenkinson 2002). This step yields, every 800 msecs, an estimate of the displacement of the brain from its initial position. This displacement is expressed in terms of translation (x, y, z) and rotation (pitch, roll, yaw). I converted these measurements into an expression of the relative displacement of the camera with respect to the corneal front surface of the right eye. To do so, I manually identified on the T1 anatomical image for each subject a voxel on the corneal front surface of the right eye. This coordinate was then projected to the corresponding location in the initial echoplanar image for each movie watching acquisition. The translation of this point in the echoplanar data over time was then derived from the 6 parameters of movement of the entire brain, and then expressed as the relative change in camera position. The vectors for camera translation were interpolated to the temporal resolution of the eye videos using a shape-preserving piecewise cubic interpolation (*interp1* in MATLAB).

It is not necessarily the case that the coordinate frame of the camera is simply a transposition of the axes of the coordinate frame of the scanner. That is, the axes of the camera coordinates may not be at right angles to those of the scanner. In exploratory analyses, I found that small rotations of the scanner coordinates did not markedly improve the match between these measures and the measure of relative camera translation obtain from the eye video. Therefore, in the results presented here, I make the simplifying assumption that the camera and scanner coordinate spaces are parallel.

The code to perform this analysis is available for download (https://github.com/gkaguirrelab/mriTOMEAnalysis).

### Derivation of eye pose from movie watching acquisitions

Each of the four movie watching acquisitions for each subject was 336 seconds in length, yielding 20,160 frames for each video. The eye features in each frame were obtained and from these the eye pose was derived. To do so, the eye features were fit with the model derived for that subject from the analysis of their gaze calibration measurements. Because the OCOL imaging arrangement provides minimal leverage over the estimation of camera depth, this parameter was locked in the derivation of eye pose. An estimate of the change in camera depth over time was available, however, from the measurement of head motion provided by the MRI data. This change in relative camera depth over time was provided to the fitting routine.

The fitting procedure yielded the radius of the aperture stop and eye rotation for each frame during movie watching for the acquisition. Eye rotation can be further converted to gaze position on the screen, although these measurements are not considered here. The fitting procedure also provided the in-camera-plane translation of the eye on each frame relative to its starting position. This estimate of eye translation was then compared to the estimate derived from the brain imaging data.

The comparison between video and MR-based measures of motion could only be performed at the video time points with a valid eye pose model fit. A video frame was retained for the analysis if:

1. A glint was present in the image frame
2. The root mean squared error of the fit of the model pupil ellipse to the pupil perimeter points was < 2 pixels.
3. The pupil perimeter points covered >75% of the circumference of the pupil ellipse

This filtering step removed video frames in which, e.g., the pupil and glint were not visible during a blink. The comparison to MR-based translation was made if, after filtering, at least 25% of the time-points of a given acquisition were still available.

## Results

I examined in empirical gaze calibration data the ability of model-based search to fit biometric parameters, and then used these parameters to recover eye pose and eye translation in previously unseen data.

### Model-based fitting of empirical gaze calibration measurements

Infrared video images of the right eye were collected from thirty participants while they performed four separate gaze calibrations (Figure 5). During each of the gaze calibration acquisitions the subject fixated sequentially upon nine targets (in random order) arranged in a grid on a display (as depicted in Figure 4a). The eye features (pupil perimeter and glint location) were then extracted for a single, representative frame from each fixation.

**Figure 5.**
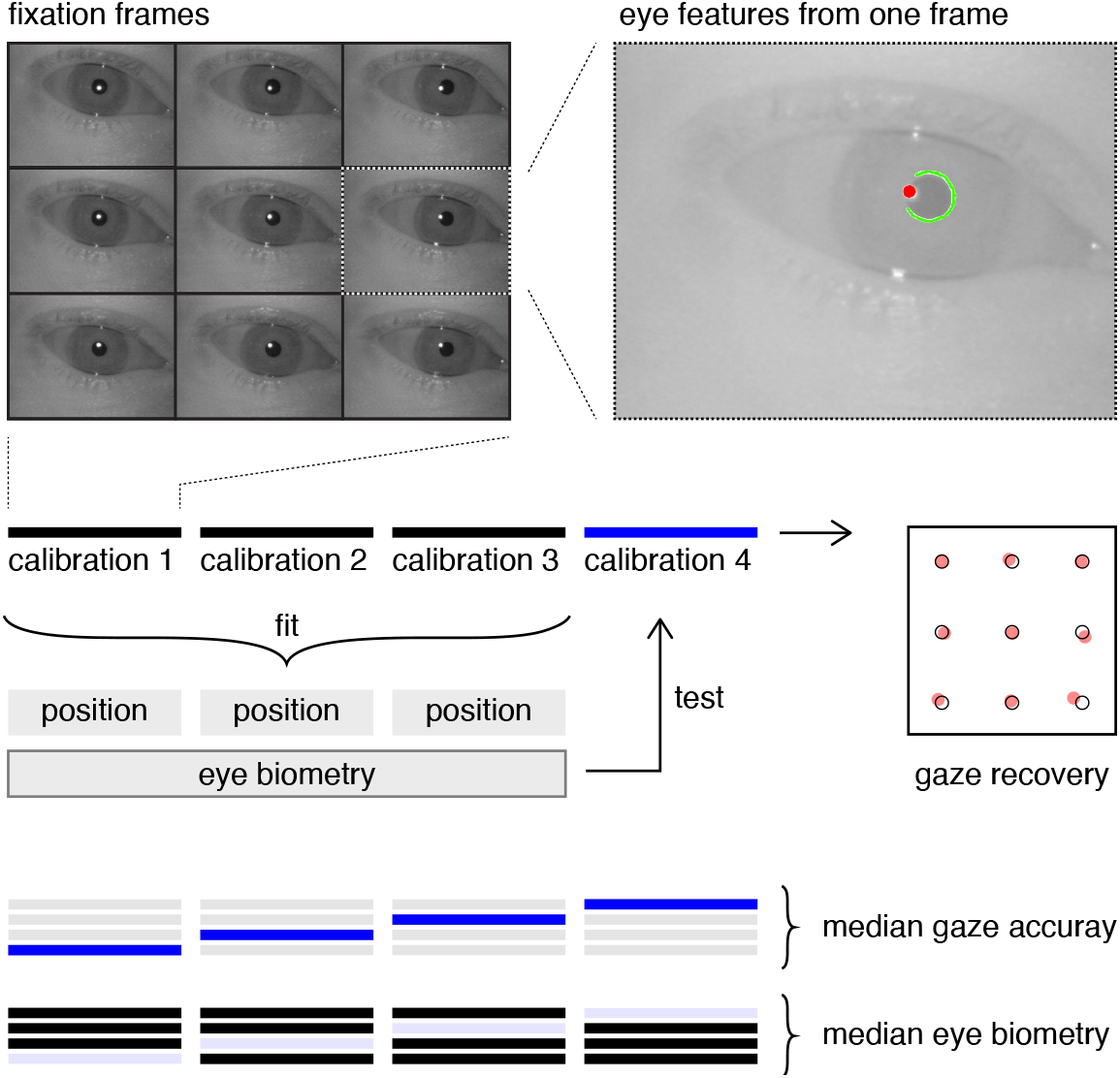
Analysis of gaze calibration acquisitions. Each subject provided data from four gaze calibration acquisitions, from which nine fixation frames were selected. The pupil and glint features were extracted from each of these frames. In a cross-validated, leave-one-out approach, the features from three of the gaze calibration acquisitions were fit in a model-based search to identify the common biometric parameters of the eye. These biometric parameters were then applied to the held-out, “test” acquisition to obtain gaze position. The recovered gaze positions were compared to the known locations of the fixation targets. The median gaze accuracy and biometry was examined for each subject across the 4-fold analysis.

A non-linear search was performed to identify the relative camera translation, eye biometric values, and eye poses that could best fit the pupil and glint features and match the rotations of the eye to the fixation targets. The measurements from three of the four gaze calibrations were fit simultaneously, with relative camera position allowed to vary between calibrations, but eye biometry locked across the three measurements. The eye biometry values obtained from this search were then used to fit the features from the fourth, held-out gaze calibration. The full set of cross-validated analyses was performed. From these analyses we can examine both the distribution and stability of the estimated biometric parameters, and the ability of the model to recover gaze position in the held-out data.

### Measurement of corneal curvature

The model fit provides an estimate of corneal curvature, which can be expressed in conventional keratometric form. For each subject, the four leave-one-out model fits yielded four estimates of k1 and k2. Figure 6a presents the median corneal curvature values obtained for each subject, along with the across-subject median (and inter-quartile range) of these measures. The obtained values are reasonably distributed. Within each subject, the four cross-fold fits support a jackknife estimate of the standard error of the measurement of corneal curvature within subject. The across-subject median of the standard error of measurement of corneal curvature was 0.63 and 0.71 diopters for k1 and k2, respectively.

**Figure 6.**
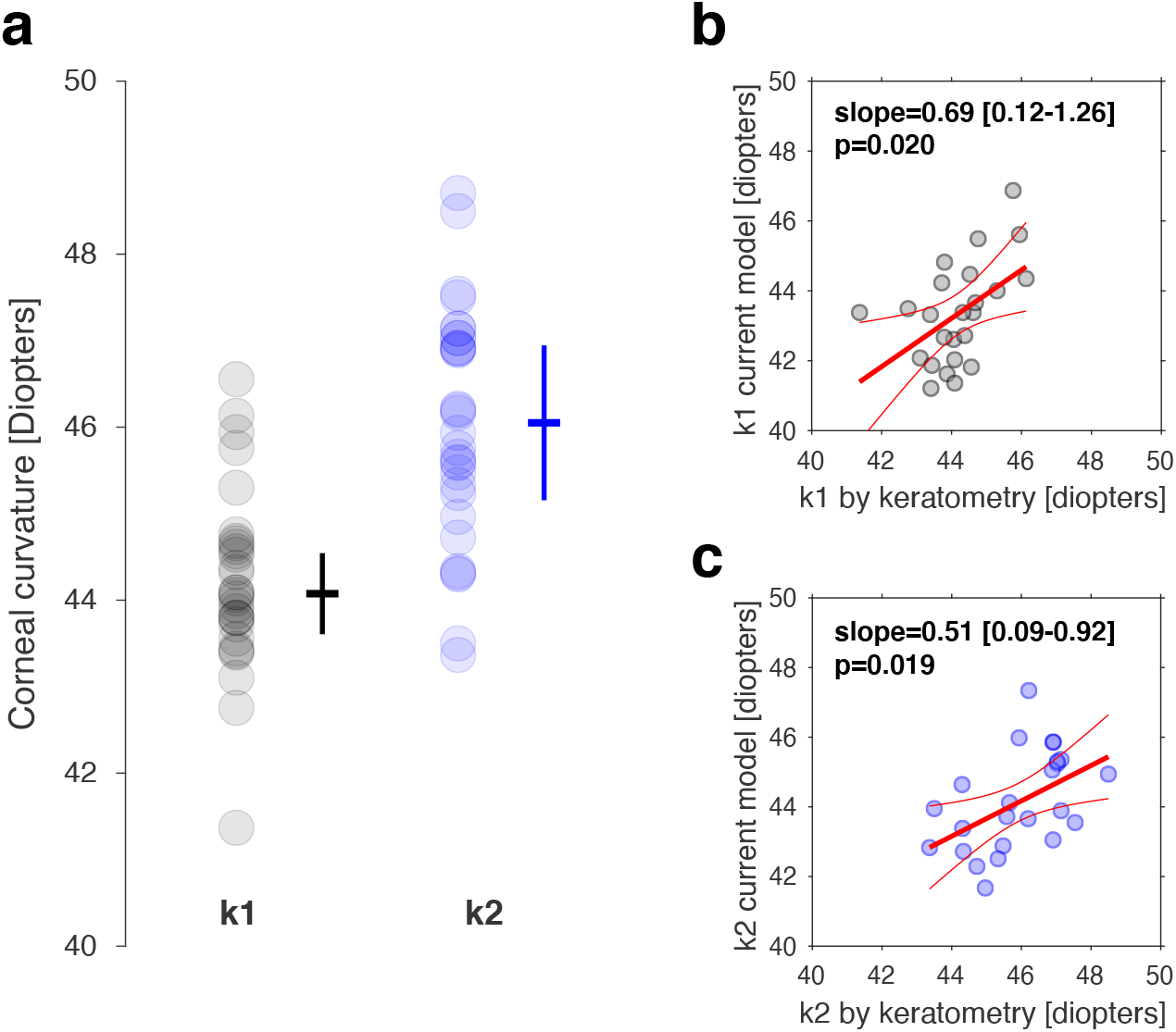
Recovery of corneal curvature. **a)** The k1 and k2 corneal curvature values obtained from each subject by model-based fitting of the gaze calibration data. The across-subject median value is indicated by the horizontal hash, and the inter-quartile range by the vertical whiskers. **b)** Twenty-four of the subjects had a separate measure of keratometry using a clinical ophthalmologic instrument. The relationship between the two measures of k1 is shown across subjects. The slope, [95% confidence interval], and p-value of a robust regression (using bi-square weighting) is shown. **c)** The relationship between the two measures of k2.

Twenty-four of the thirty subjects also had a measurement of corneal curvature provided by clinical keratometry. Figures 6b and 6c examine the agreement across subjects between the measure of corneal curvature provided by a clinical keratometric device, and by the current model fitting approach of the pupil and glint appearance. The current model provides an estimate of k1 that is unbiased as compared to the keratometric standard, and the 95% confidence interval of the across-subject slope of the relationship includes unity. For k2, the current approach yields values that somewhat under-estimate the measure provided by keratometry, although there is still a significant, positive relationship between the measures. Overall, the ability of the current approach to recover corneal curvature is good, in particular given that this estimate is based only on the shape and location of the pupil and first Purkinje image, and was performed on subjects for whom 1/3^rd^ were wearing corrective lenses during the measurement.

The model fit includes parameters for the rotation of the corneal surface. For most subjects (26/30), the estimate of corneal torsion was consistent with the steep meridian of the cornea being oriented vertically (80-100°), as is most commonly seen in normally sighted populations (Tonn 2015). Four subjects had values consistent with oblique astigmatism (angles of ~120° or ~70°). In three of these subjects, however, the jackknife estimate of the standard error of the measurement was large (>10°). These subjects also had relatively little astigmatism (the difference between k2 and k1 was <2 diopters). When corneal astigmatism is small, and the cornea front surface approaches rotational symmetry, the effect of corneal torsion upon the model output shrinks and the estimation of this parameter would be expected to be unstable. For one subject, who had astigmatism >3 diopters, an oblique corneal torsion was found with a small (5°) standard error. This finding suggests that the model search is able to measure corneal torsion under suitable conditions, although there were very few subjects in the current study who allowed for a full examination of this performance. For the other two parameters of corneal rotation (“tilt” and “tip”) there was minimal variation across subjects or departure from the initial values supplied to the model.

### Measurement of rotation depth

The model fit includes parameters that control the depth of the center of rotation of the eye, with separate rotation centers for horizontal and vertical eye movements. Figure 7a presents the median observed depth of the rotation center of the eye for horizontal and vertical eye movements for each of the 30 subjects, as well as the median, across subject values. A clear aspect of these data is that the across-subject rotation center depth for horizontal eye movements (13.90 mm ± 0.33 interquartile range) is reliably and substantially larger than for vertical eye movements (12.23 mm ± 0.34). These measurements are quite similar to prior findings (Fry & Hill 1962, 1963; Hayami 2002). Across subjects, the rotation depths estimated for horizontal and vertical movements were positively correlated (Pearson’s r = 0.68).

**Figure 7.**
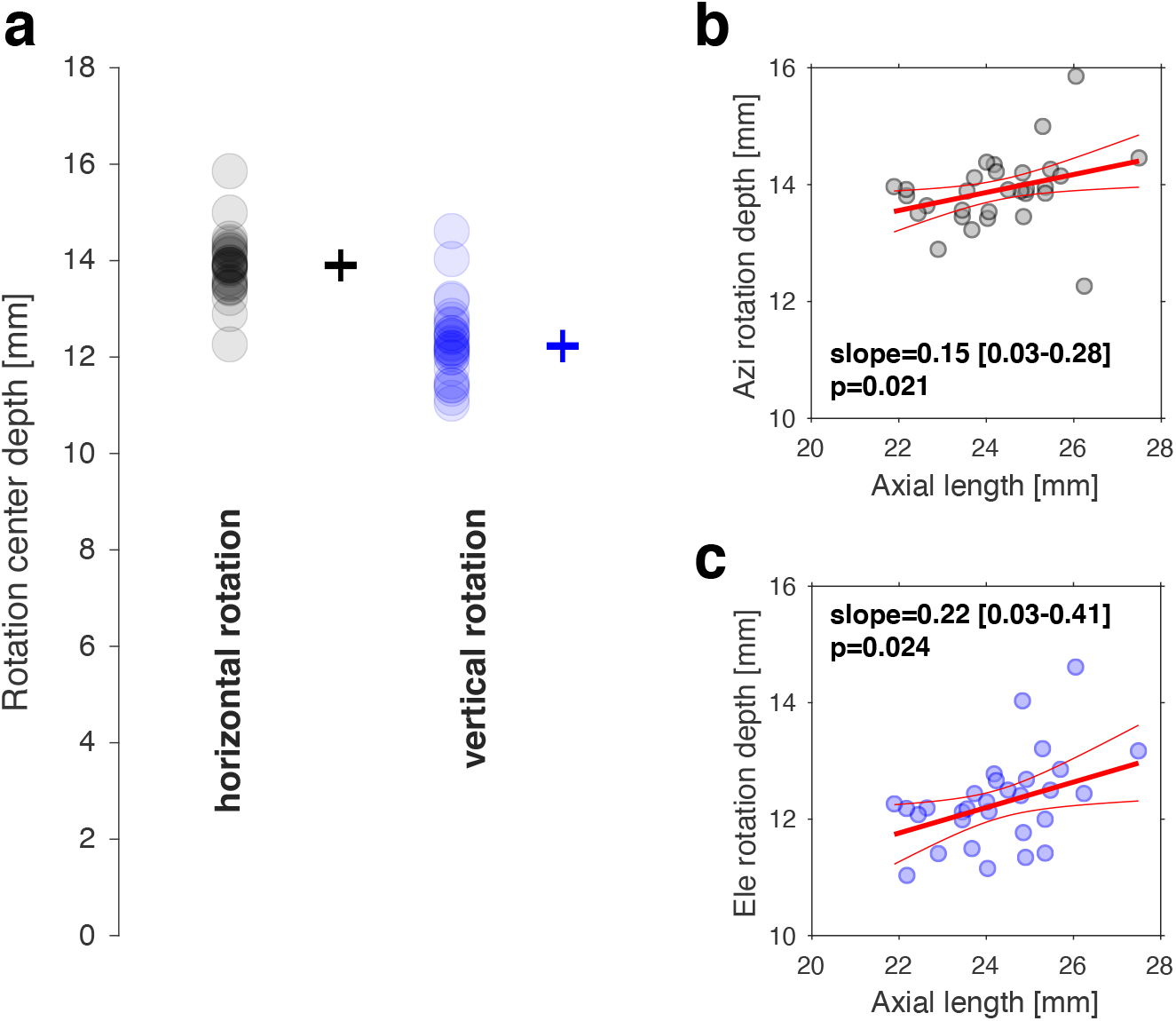
Measurement of rotation center depth. **a)** The depth of the horizontal and vertical rotation centers were measured for each subject by model-based fitting of the gaze calibration data. The across-subject median value is indicated by the horizontal hash, and the inter-quartile range by the vertical whiskers. **b)** A measurement of the axial length of the eye by a clinical ophthalmologic instrument was available for each subject. The relationship between axial length and horizontal rotation center depth is shown across subjects. The slope, [95% confidence interval], and p-value of a robust regression (using bi-square weighting) is shown. **c)** The relationship between axial length and depth of the vertical rotation center.

As was examined for corneal curvature, the cross-fold model fits performed in each subject support a jackknife estimate of the within-subject standard error of the measurement of the rotation centers. The across-subject median of the standard error of measurement of the depth of the horizontal and vertical rotation centers was 0.29 and 0.28 mm. This small within-subject error given the ~4 mm range of depth values across subjects suggests that there are reliable individual differences in this biometric property. Variation in axial length has been found to be correlated with the depth of the horizontal center of eye rotation (Dick 1990). We examined in our data the relationship between measured axial length in each subject and the depth of the horizontal and vertical centers of rotation (Figure 7b, c). A significant association between axial length and rotation depth was found for both measures. The slope of the relationship was ~0.2, implying that for every 1 mm increase in the axial length of the eye, rotation center depth increases by 0.2 mm.

### Measurement of gaze position

These empirical findings indicate that the model-based search is able to recover meaningful biometric parameters for each eye under study. The simulations presented in Figure 3 suggest that, given accurate biometric parameters, model-based search should be able to return the pose of the eye, and from this a measure of gaze position. For each of the cross-validated measurements, the biometric parameters derived using three of the gaze calibration acquisitions were used to predict gaze position for the held-out, fourth acquisition. From each held-out acquisition I measured the median, absolute error in recovering the visual field positions of the nine fixation targets, and for each subject I identified the median such performance across the four held-out tests. Overall, across all targets, acquisitions, and subjects, the model-based search was able to recover gaze position in previously unseen data with a median absolute error of 0.58°.

Figure 8a presents the nine fixation locations predicted in the acquisition with the median prediction error in each subject. The agreement between the veridical target locations and the predicted gaze positions is evident. Figure 8b presents a histogram of the median prediction error across targets and acquisitions for each subject. For 28/30 subjects, the median absolute prediction error was less than 1°.

**Figure 8.**
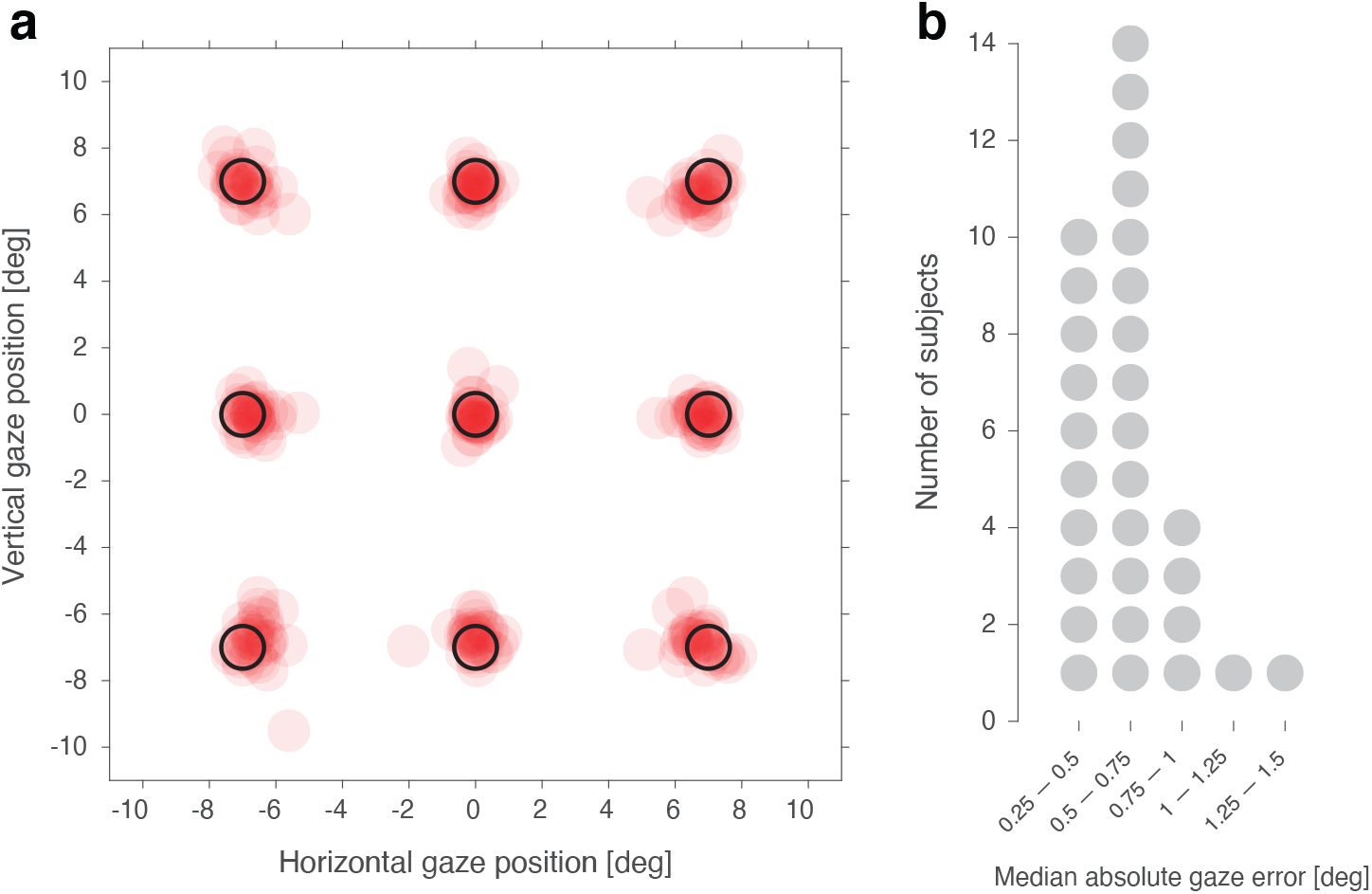
Prediction of gaze location. **a)** The biometric parameters derived from three gaze calibration acquisitions were used to predict the gaze locations in the fourth, held-out acquisition. The median-performing test was selected for each subject. Shown are the predicted gaze locations (in red) for each subject, along with the veridical target locations (in black). **b)** The distribution of median absolute gaze errors across the thirty subjects.

The quality of gaze prediction in the held out acquisition was strongly related to the quality of the model fit to the training acquisitions. The search to derive biometric parameters attempts to minimize error in matching eye rotations to the fixation targets. The Pearson’s correlation between this gaze error in the initial model fitting, and the gaze error in the held-out acquisition, was 0.87. Two subjects had gaze prediction errors >1°. For one of these subjects there was no apparent pattern to the gaze prediction errors, and inspection of the video recordings suggests that poor fixation stability was the cause of the high error. For the second subject, the modeled rotation of the eye consistently under-predicted the position of the target in degrees of visual angle. While this subject was myopic (–6.25 D), and wore spectacle lenses during data collection, neither of these factors appear sufficient to explain the poor model fit. Several other subjects had greater myopia (up to –10.25 diopters), and wore artificial lenses, but did not have this pattern of poor fit of the model. I consider in the discussion possible explanations for the poor model fit in this subject.

### Measurement of eye translation

Analysis of the gaze calibrations provided biometric parameters for each subject. These parameters were then used to perform model-based fitting of video acquisitions obtained from each subject while they watched a movie. This analysis provided a continuous measure of the pose of the eye (pupil size and rotation angles), as well as the relative displacement of the camera from its initial position, over the course of the movie-watching acquisition. Supplementary Movie 3 shows the fit of the model to an example eye video. Here I examine the agreement of the measurement of camera translation with an independent measure provided by observation of the position of the head over time.

Recording of the movie-watching eye video took place during a functional MRI scan, providing a simultaneous, volumetric measurement of the brain every 800 msecs. Analysis of the functional MRI data yielded a measurement of the rigid-body displacement of the brain from its initial position over time. This motion measure consists of three directions of translation and three dimensions of rotation of the brain in the coordinate frame of the MRI scanner (Figure 9a, left). In the example measurement of head movement from one acquisition (Figure 9b) a slow rotation and translation of the head towards the subject’s left can be seen over the course of the 336 second recording. If we select a single point on the corneal surface of the right eye, then the six parameters of head motion may be described instead as three parameters of translation in camera coordinates (Figure 9a, right). The combined effect of a yaw rotation and translation of the head is a lateral translation of the eye by about 2 mm over the course of the measurement (Figure 9c).

**Figure 9.**
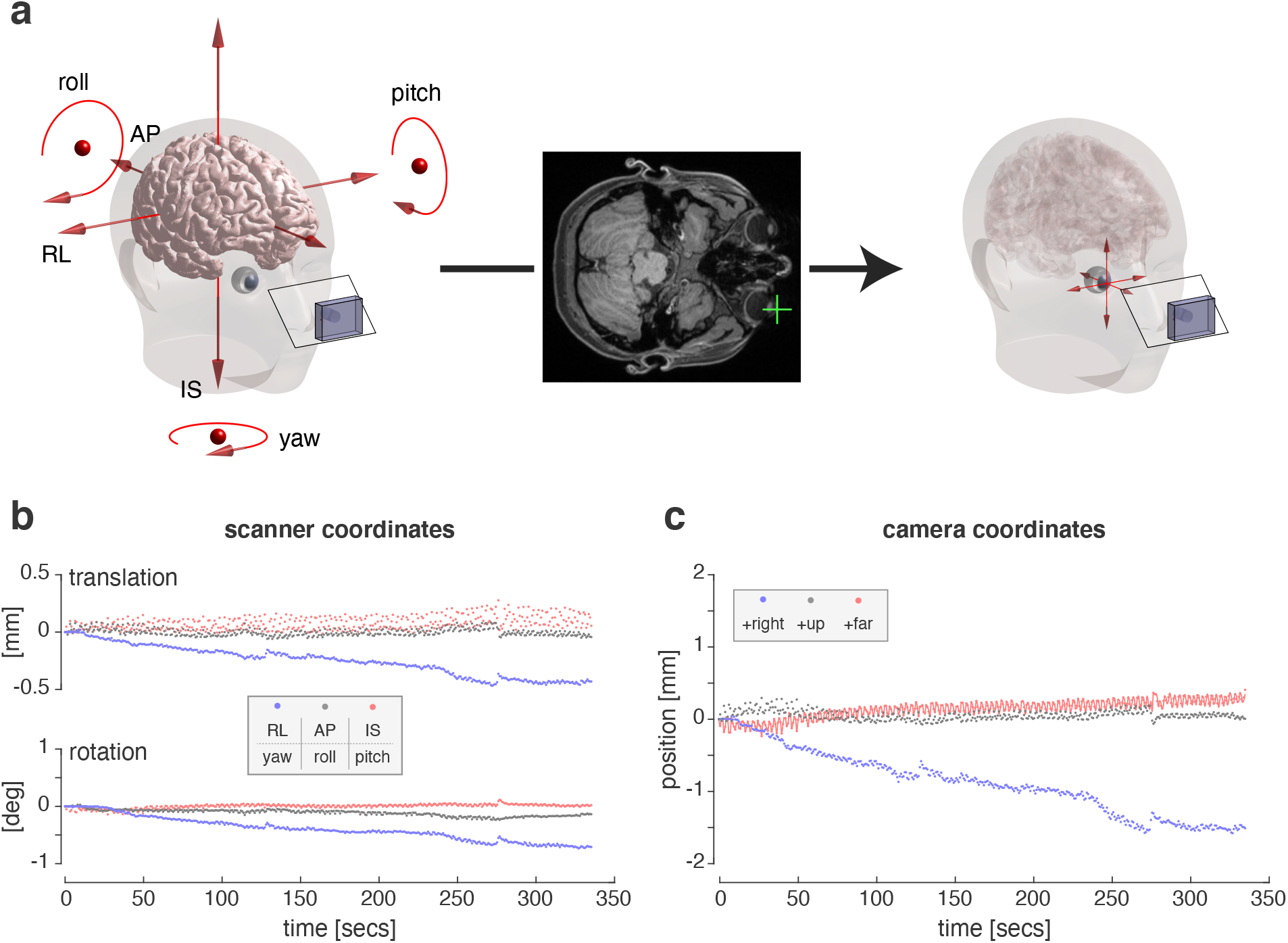
Derivation of relative camera movement from MRI data. **a)** The geometric arrangement of the subject’s head, brain, and eye are shown, along with the video camera and an angled cold mirror (left). The functional MRI data provides an estimate of the motion of the brain over time in terms of six degrees of freedom. RL = right-left, AP = anterior-posterior, IS = inferior-superior. The anterior-most point on the cornea of the right eye was identified in a T1-weighted MR image for each subject (center). The brain motion vectors were converted into an expression of relative camera motion with reference to this point on the cornea (right). **b)** The head motion values measured by MRI during a movie-watching acquisition for one example subject. **c)** The predicted relative translation in camera image coordinates for the right anterior corneal surface. The values for the depth vector have been fit with a smooth interpolation (red line).

One may have the concern that reducing the 6 parameters of movement of the head to only the translation of the anterior surface of the eye discards important information regarding head rotation. In the current model, eye rotation is defined with respect to the optical axis of the camera. The camera and screen coordinate systems are fixed with respect to one another. We may presume that head rotation is accompanied by a counter-rotation of the eyes so that the subject continues to fixate upon the screen. Under these circumstances, a pure head rotation (in the absence of eye translation) has minimal effect upon the eye pose and camera translation values derived from the model fit to the video images. Only the value for the primary position of the eye (which is defined here in relationship to the optical axis of the camera) would be altered by head rotation.

We may compare the estimate of relative camera translation obtained from brain imaging to that provided by the model fit to the movie-watching eye video. The one-camera, one-light source approach used to record videos of the eye, however, is poorly suited to measurement of changes in relative camera depth (Figure 3, bottom row). Therefore, the change in camera depth estimated from the head motion measurement was supplied to the model used to fit eye pose, after up-sampling the depth estimate to the temporal resolution of the video (Figure 9c, red line). We may then compare in-camera-plane translation measured from the MRI scan to that measured by model fit to the eye video.

Figure 10a shows the relative, in-image-plane camera translation for the example acquisition as measured by the MRI scanner and the model fit to the eye video. There is good agreement between the measurements. This clear for the relative horizontal translation of the camera, in which the evolution of a 2 mm shift in position is well correlated between the MRI-based and video-based measurements (Pearson’s r=0.97). The absolute range of motion is smaller in the vertical direction, especially for the MRI-based measurement. The video measurement has a notable, high-frequency oscillation, with a cycle time (13/minute) suggestive of the respiratory cycle. The lower correlation (Pearson’s r=0.62) of the measurements of vertical camera translation seem to reflect that the coarse temporal sampling of the MR imaging is less sensitive to this motion component.

**Figure 10.**
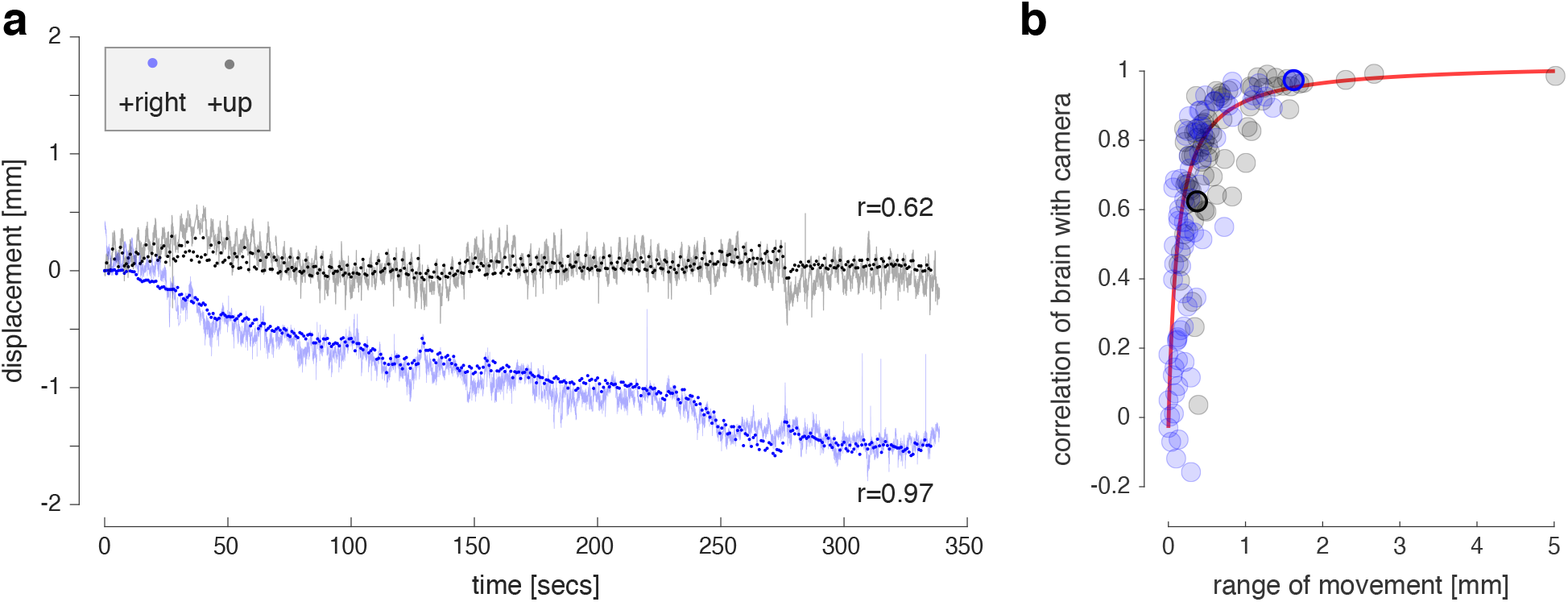
Comparison of eye translation as derived from MR and model-based fitting of video images. **a)** The in-image-plane horizontal (blue) and vertical (black) translation of the eye is estimated from MR data (dots) and from fitting the simultaneously acquired eye video (transparent lines). The Pearson correlations between the measures is given. **b)** The correlation of MRI and video-based measures of head motion for horizontal (blue) and vertical (black) translation, as a function of the maximum displacement present in the MRI data for that acquisition. The plot symbols associated with the plot shown in panel (a) are marked with a thicker outline. The red line is the fit of a 3-parameter reciprocal function.

A total of 78 movie-watching acquisitions were available for analysis, each yielding a correlation between video and MRI-based measurements of camera translation in the vertical and horizontal direction. I considered the possibility that this correlation would be reduced in the setting of minimal head motion. Figure 10b plots the correlation between MRI and eye video translation measures for each of the movie watching acquisitions as a function of the range of motion present in the MR measure. When there is at least 0.5 mm of head motion, the correlation between the two measures is quite high (Pearson’s r>0.75). Below this threshold, however, the two measures diverge. It is not clear if the disagreement between the measures below this threshold of movement is due to inaccuracy of the model-based search, or due to the relative insensitivity of the MRI-based measurement for small, higher temporal frequency motion. Nonetheless, we can at least confirm that the model-based search recovers relative camera translation with a precision of less than 1 mm.

## Discussion

I have presented here a software model of the appearance of the human eye as it undergoes biologically accurate rotation, with a particular focus upon the elliptical form of the entrance pupil and the location of the first Purkinje image (the “glint”). In simulations and empirical measurements, I show that a model-based fit to these features may be used to recover the ellipsoidal form of the cornea, the depth of the rotation centers, the rotation pose of the eye, the size of the aperture stop, and relative camera translation. I take these findings as evidence of both the accuracy and utility of the model.

### A general model of eye appearance

There is an extensive prior literature that concerns the derivation of gaze location based upon the appearance of the eye. The current study falls within the category of feature-based modeling approaches. Generally, the shape and location of one or more segmented features of the eye are measured—such as the pupil, glint, or outline of the iris—and are then related to gaze position via a set of equations based upon a simplified eye model (Hansen 2010). These equations are specific to the particular eye features or imaging approach used. For example, different solutions are required based upon the number of cameras or light sources, or if the measure is monocular or binocular.

The current study adopts a different approach. Instead of defining a fixed set of equations, I examine the performance of fully model-based search to derive eye pose and translation. Fitting of empirical data begins with a ray-traced, parametric model of the human eye. The output of the model, given a set of candidate parameters, is used to simulate the appearance of the features of the eye, and the model parameters are adjusted to achieve the best agreement between the simulation and the empirical measurements. A key advantage of this forward-model approach is that it is easily generalized. For example, the model may be interrogated to describe the appearance of different eye features, as viewed with multiple cameras or light sources.

The ray-tracing simulation can provide the appearance of the eye as observed through artificial lenses worn by the subject to correct spherical ametropia. Three effects of corrective lenses are captured in the current simulations. First, a spectacle or contact lens refracts the image of the eye as seen by the camera. Second, a contact lens alters the curvature of the tear film, changing the behavior of the glint. Third, a spectacle lens introduces magnification or minification of the world as seen by the eye, altering the visual angle at which gaze calibration targets are presented and thus the relationship between eye rotation and the position of stimuli on a screen. These subtle and interacting aspects of eye appearance are readily handled by a model-based fitting approach.

There have been recent advances in the development of appearance-based gaze tracking routines. In these approaches, a machine learning (or similar) algorithm is used to learn the mapping between the image appearance of the eye and face and the location of a target upon which the subject is fixating (Wang 2016; Yiu 2019). Such efforts benefit from a rich supply of training images that capture biological and environmental variability. Świrski and Dodgson (2014) proposed the use of computer graphics rendering software to create synthetic ground-truth images to evaluate and train gaze tracking approaches. The current model may find similar application, in particular for feature-based tracking techniques.

The software I have developed includes two search routines that support fitting the model to empirical measurements of eye appearance. Each is a non-linear search across a set of parameters that define the appearance of the simulated eye features. The core, “eye pose” search identifies the pose and translation of an eye that best accounts for the location of image points that are on the border of the entrance pupil, or at the center of a glint. The eye pose search then serves as the core of a second, “biometry” search routine that estimates parameters of the eye model by matching rotation angles of the eye with the visual angles of a set of targets. The particular properties of these routines, including the composition of the objective function and the boundaries and staging of the parameter search, were arrived upon after working with the empirical datasets studied here. It may be the case that changes to the search routine would be desired for different datasets. It should also be noted that these routines are computationally intensive. For example, an eye pose search might require 250 iterations of the ray-tracing simulation to converge upon a result, taking 1-2 seconds to perform on a standard laptop CPU.

### Recovered eye pose, translation, and biometric values

Video measurements of the eye are often used to estimate dynamic and static properties of the subject under study. In the current study I examined the accuracy of several of these estimated parameters in empirical data.

In many cases, the goal of video based eye tracking is to estimate the point of gaze of a subject who is being observed, although there is considerable variability in the manner in which the accuracy of this estimate is evaluated (Kar 2017). I adopted here a fully cross-validated approach. The biometric parameters of the model were fit using the features from a set of eye images, and then the model was used to estimate gaze locations in a held-out dataset from that subject. Across subjects, the predicted gaze locations agreed with the fixation target locations with an absolute median error of 0.58°. This error incorporates any instability on the part of the subject in maintaining fixation upon a target, which itself is on the order of 0.3° standard deviation (as measured by scanning laser ophthalmoscopy; Rohrschneider 1995). The accuracy of the current approach, in particular for a one camera, once light source system in the presence of a small degree of head motion, is quite good compared to other techniques (Kar 2017).

The eye pose search provides as well the radius of the aperture stop of the iris. Notably, the model is expressed in terms of the physical aperture of the iris, which then gives rise to the entrance pupil as seen through the refractive effects of the cornea. I do not have available an independent measure of this physical property of the eye to which I can compare the model output. As I have previously shown, however, the model output provides an accurate account of the shape of the entrance pupil (Aguirre 2019). Given that the physical and optical properties of the cornea are modeled based upon empirical measures of these structures, there is reason to expect the model-based estimation of the size of the aperture stop is also accurate.

Movement of the head with respect to the camera complicates eye tracking approaches. One class of solution is to use a head-mounted camera (e.g., Dierkes 2019). Other techniques attempt to reduce the effect of head translation upon the estimation of gaze location. In the current approach, the translation of the camera relative to the head is obtained as part of the model-based search. In simulations I show that translation within the camera image plane is accurately measured using a one-camera, one-light imaging system. A notable feature of the current work is that the accuracy of the estimate of head translation is validated by comparison to a separate measure obtained by simultaneous MR imaging of head. I find that these measures agree to a precision of 0.5 mm.

Finally, fitting of gaze calibration data with the current approach provides estimates of the rotation centers of the eye and the optical power of the cornea. In empirical data, I find clear evidence for the existence of separate rotation centers for horizontal and vertical eye movements, replicating the original findings of Fry and Hill (1962 and 1963). I also replicate the prior observation that the depth of the center of rotation varies with the axial length of the eye (Dick 1990). After accounting for this effect of axial length, the measurements reported here indicate that, in the average emmetropic eye, the mean horizontal center of rotation is at a depth of 13.8 mm, and the vertical center of rotation is at a depth of 12.1 mm. The parameters of the model also include the curvature of the astigmatic cornea, and the model parameters derived here agree well with the direct measure of corneal curvature provided by a clinical instrument.

I am not aware of a prior approach to the analysis of eye tracking data that recovers this entire set of parameters simultaneously, nor does so with this degree of validated accuracy.

### Modelling assumptions and a poorly fit subject

The model was unable to account for the eye movement of one of the thirty subjects. For this subject the modeled rotation of the eye was consistently less than would be predicted to bring a point in visual space into fixation. In their 1962 study, Fry and Hill observed that three of their thirty-one subjects had eye movements that did not correspond to a pure rotation, but included as well an element of translation. The approach that I have developed here incorporates assumptions regarding eye movement and fixation that would lead to poor fitting of gaze position for such subjects. In the model-based search to fit gaze calibration measurements, a change in the position of the eye is assumed to be a pure rotation about the vertical and horizontal rotation centers. While the eye-pose search is able to measure relative translation of the eye, the fit to gaze calibration data attempts to find parameters of the model that minimize the measured eye translation. While not explored here, it would be possible to use the current model in a modified search approach that attempted to measure eye translation with changes in fixation. Such an attempt would benefit from data in which head motion is strictly minimized by (e.g.) use of a bite bar.

The search algorithms I have studied here also assume the equivalence of the optical and rotation centers of the eye, which is certainly not the case. A refinement of the search to remove this assumption could be implemented using the current model eye. The model supports ray-tracing through the optics of the eye to the retina, with routines for calculation of the line-of-sight and optical center of the eye (Aguirre 2019; Chen 2020). This additional modeling effort would be desirable for circumstances in which the effect of ocular parallax is expected to be noticeable, such as when fixating upon close targets.

### Depth and Torsion

Two aspects of eye position were difficult to measure in the empirical data examined here. First, as demonstrated in simulation and consistent with prior results, the distance of the camera from the eye of the subject is poorly measured with the one-camera, one-light source imaging system used to collect the empirical data. To allow the modeling to proceed, the initial distance of the camera from the eye of each subject was set by hand based upon image appearance. This distance estimate was then updated in the movie-watching data by deriving the change in camera depth from the MR images of the head. As demonstrated in simulation, the current model fitting approach can readily recover camera depth information if multiple light sources (or cameras) are used.

The current model has several parameters that relate to the torsional rotation of modeled components about their optical axes. The camera, astigmatic corneal front surface, and eye all have independent torsional parameters. Further, the dynamic rotation of the eye may include both “true” torsion, as well as a corrective, “pseudo” torsion to allow the simulation to follow Listing’s Law. This complexity, however, found little application in the empirical data studied here, as infra-red images of the eye provide minimal leverage upon the relative torsional rotation of camera, cornea, and eye. In the absence of iris features or other marks of polar position around the eye, torsional information is provided only by the slightly elliptical shape of the aperture stop of the iris, the astigmatism of the cornea, and the inequality of the horizontal and vertical centers of rotation. Similar to camera depth, camera torsion was set by hand for analysis of the empirical data, in this case based upon the measured canthal angle. Beyond this, I performed only a limited examination of torsional parameters in the current study. Nonetheless, the software framework is available within the current model to derive torsional parameters given appropriate eye features.

## Conclusions

The current work is in the spirit of other efforts in vision science to create forward models that incorporate accurate, quantitative descriptions of the underlying biology (e.g., Cottaris 2019). I have focused here upon one particular application of the model eye, namely tracking the gaze of an observed subject. The software model is open source, well commented, and contains numerous examples. My hope is that this code base will support further studies to understand and measure the human eye in motion.

## Acknowledgments

Thank you to Giulia Frazzetta, Edda “Briana” Haggerty, Saguna Malhotra, and Laura Cutler for their assistance in setting parameters for the extraction of eye features from the raw videos. Thank you as well to Yu You Jiang and Jessica Morgan for their kindly sharing the clinical axial length and corneal curvature measurements for the studied subjects. Nicolas Cottaris provided helpful feedback on an initial draft of the paper. Figure 9 includes the 3D renderings “Human Brain, Full Scale” by “MiloMiis”, and “Human Head (NURBS)” by “grozny”, both made available under the Creative Commons, Share Alike license, and downloaded from www.thingverse.com. This work was supported by National Institutes of Health Grant U01EY025864: Human Connectomes in Low Vision, and P30 EY001583: Core Grant for Vision Research.

## Supplementary methods

### Modeling of artificial lenses

Spectacle and contact lenses are defined in the model by their optical power in diopters. For spectacle lenses, the lens material was modeled with the refractive properties of polycarbonate. A lens vertex distance of 12 mm was assumed. A simulated “grind” was used to produce an ophthalmic lens (also known as a meniscus, or “convex-concave” lens) of the desired optical power. The base curvature (front surface power) was set using Vogel’s rule. The position of the front surface and the curvature of the back surface was then found by a non-linear search which attempted to match the desired optical power with that measured via ray-tracing through the candidate lens. This search was subject to non-linear constraints to create lenses of the appropriate shape and thickness (e.g., to enforce a minimum thickness at the lens center for a minus lens).

A similar procedure was followed for the specification of contact lenses. The index of refraction of hydrogel was assumed, and the back surface of the lens was set to match the curvature of the front surface of the cornea.

Artificial lenses produce magnification / minification of the visual world from the perspective of the lens wearer. For the current work, spectacle magnification is relevant for the modeling of gaze position. The effect of artificial lenses upon angular magnification as seen by the retina was calculated by ray-tracing simulation. To do so, a set of world targets were modeled and the effect of the artificial lens upon the angle of rays intersecting the center of the entrance pupil was measured. The accuracy of the simulation was evaluated by comparing the output of the model to calculations reported in Figure 1 of Westheimer (1963). Figure S1 shows the percentage of magnification (positive values) or minification (negative values) produced by spectacle and contact lenses of varying optical power. Matching Westheimer, this calculation was performed for an emmetropic eye with a corneal curvature of 43.5 keratometric diopters and without corneal astigmatism. The current result is in good agreement with Westheimer.

**Figure S1.**
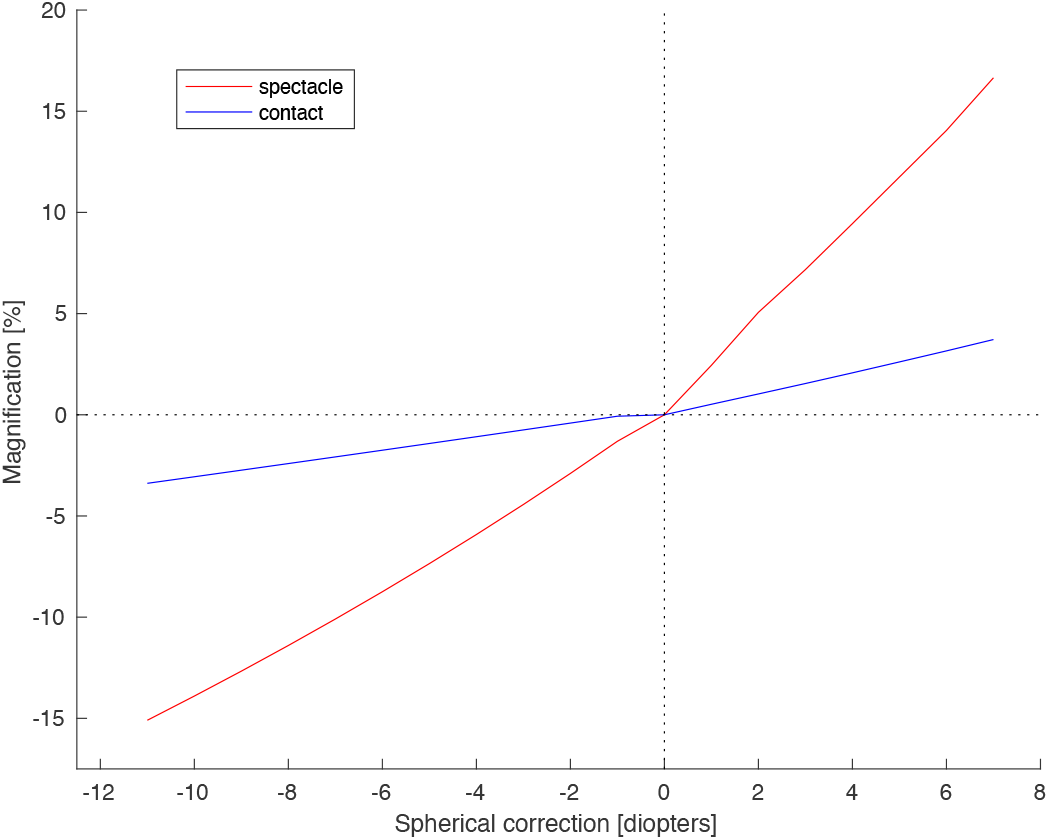
Measurement of the magnification produced by corrective lenses. Compare with Figure 1 of Westheimer 1963.

The contact lens surfaces rotate along with the rest of the eye, while the spectacle lens remains at a fixed position with respect to the camera. Spectacle magnification therefore induces a change in the magnitude of eye rotation needed to fixate a target. To account for this effect, the modeled visual position of the gaze calibration targets was scaled by the spectacle magnification calculated for any subject who wore spectacles during the measurement.

### Supplementary movies

**Movie S1**. Illustration of model-based fitting of eye pose and camera translation. A default set of biometric parameters was used to generate a simulation of the appearance of the entrance pupil and the glint of an eye observed by a camera. The particular eye pose and camera translation values used for this simulation are given as green text at the bottom of frame. Ten points on the perimeter of the entrance pupil (black closed circles) and the location of the glint (red open circle) were defined. Model-based search was then used to identify camera pose and eye translation values that would best account for the pupil and glint data points. For each iteration of the non-linear search, the model eye is rendered using parameters of the search given in black text. Shown are the predicted pupil ellipse (green) and glint location (red asterisk). The cornea (yellow) and vitreous chamber (yellow) are illustrated as well. Over iterations of the search, the progressive refinement of the parameters steadily improves the fit of the simulated data. At termination, the search has recovered parameters that are extremely close to those used to generate the simulated data.

**Movie S2**. Illustration of model-based fitting of gaze calibration data. The search across biometric parameters to fit a set of gaze calibration measurements is shown. Nine frames from a gaze calibration video are shown in a grid on the left, with each frame obtained while the subject was fixating upon a different target location. As the search for biometric parameters proceeds, a render of the candidate model eye is superimposed on each of the video frames. The red line indicates the plane of horizontal eye rotation. Presented on the right are parameters and outputs of the search. At top is shown the match between the gaze target locations (black circles) and the gaze locations predicted from the rotations of the eye returned from fitting the eye features with the candidate eye model. In the middle are the set of biometric and scene parameter values at each iteration; parameters that are being changed on this iteration are shown in red. At bottom is a plot of the relative error in each of the four metrics of the fit. These are 1) fit of the predicted pupil ellipse to the observed pupil perimeter points; 2) fit of the predicted glint location to the observed glint location; 3) match of the predicted gaze locations to the fixation target locations; 4) the size of individual variation in camera location between the nine different frames.

**Movie S3**. Fit of the model to a continuous eye recording. Shown is an infra-red video recording at 60 Hz of a participant who was watching a movie. For each frame of the video, points were found on the border of the entrance pupil (white dots) and in the center of the glint (red dot). These eye features were then fit on each frame with a model-based search. The model made used of biometric parameters for this subject, derived from gaze calibration measurements. The posed model eye is superimposed on each video frame for which the model was able to achieve a good fit of the data; note that the model is absent when the subject blinks. Over the course of the 5 minute recording, the eye translates towards the subject’s left; this can be seen clearly by scrubbing through the video at high speed. Note that the fit to the data is sensitive to this change in relative camera position, and translates the model eye accordingly.

## References

Acerbi, L. & Ma, W. J. (2017). Practical Bayesian Optimization for Model Fitting with Bayesian Adaptive Direct Search. In Advances in Neural Information Processing Systems 30, pages 1834–1844.

Aguirre, Geoffrey K. “A model of the entrance pupil of the human eye.” Scientific reports 9.1 (2019): 110.

Atchison, David A., and George Smith. Optics of the human eye. Butterworth-Heinemann, 2000.

Bingham, Geoffrey P. “Optical flow from eye movement with head immobilized:”Ocular occlusion” beyond the nose.” Vision Research 33.5-6 (1993): 777–789.

Bouguet, J. Y. “Camera Calibration Toolbox for Matlab.” Computational Vision at the California Institute of Technology. 2012.

Chen, Min, et al. “The influence of axial length upon the retinal ganglion cell layer of the human eye.” Translational vision science & technology 9.13 (2020): 9–9.

Nicolas P. Cottaris, Haomiao Jiang, Xiaomao Ding, Brian A. Wandell, David H. Brainard; A computational-observer model of spatial contrast sensitivity: Effects of wave-front-based optics, conemosaic structure, and inference engine. Journal of Vision 2019;19(4):8. doi: 10.1167/19.4.8.

Dick, Graham L., Bryan T. Smith, and Peter L. Spanos. “Axial length and radius of rotation of the eye.” Clinical and Experimental Optometry 73.2 (1990): 43–50.

Dierkes, Kai, Moritz Kassner, and Andreas Bulling. “A fast approach to refraction-aware eye-model fitting and gaze prediction.” Proceedings of the 11th ACM Symposium on Eye Tracking Research & Applications. 2019.

Fry, G. A., and W. W. Hill. “The center of rotation of the eye.” Optometry and Vision Science 39.11 (1962): 581–595.

Fry, Glenn A., and W. W. Hill. “The mechanics of elevating the eye.” Optometry and Vision Science 40.12 (1963): 707–716.

Fuhl, Wolfgang, et al. “Pupil detection for head-mounted eye tracking in the wild: an evaluation of the state of the art.” Machine Vision and Applications 27.8 (2016): 1275–1288.

Glasser, M. F., Sotiropoulos, S. N., Wilson, J. A., Coalson, T. S., Fischl, B., Andersson, J. L., Xu, J., Jbabdi, S., Webster, M., Polimeni, J. R., Van Essen, D. C., Jenkinson, M., & Consortium, W. U.-M. H. (2013). The minimal preprocessing pipelines for the Human Connectome Project. Neuroimage, 80, 105–124

Guestrin, Elias Daniel, and Moshe Eizenman. “General theory of remote gaze estimation using the pupil center and corneal reflections.” IEEE Transactions on biomedical engineering53.6 (2006): 1124–1133.

Gunter K. von Noorden, MD; Emilio C. Campos “Binocular Vision and Ocular Motility Theory and Management of Strabismus” American Orthoptic Journal 51.1 (2001): 161–162.

Hansen, Dan Witzner, and Qiang Ji. “In the eye of the beholder: A survey of models for eyes and gaze.” IEEE transactions on pattern analysis and machine intelligence 32.3 (2009): 478–500.

Harris, W. F. “Nodes and nodal points and lines in eyes and other optical systems.” Ophthalmic and Physiological Optics 30.1 (2010): 24–42.

Haslwanter, Thomas. “Mathematics of three-dimensional eye rotations.” Vision research 35.12 (1995): 1727–1739.

Hayami, Takehito, Kazunori Shidoji, and Katsuya Matsunaga. “An ellipsoidal trajectory model for measuring the line of sight.” Vision research 42.19 (2002): 2287–2293.

Jenkinson, M. (2002). Improved Optimization for the Robust and Accurate Linear Registration and Motion Correction of Brain Images. NeuroImage, 17(2):825, 825–841.

Nakayama, K., K. Ciuffreda, and C. Schor. “Kinematics of normal and strabismic eyes.” Basic and Clinical Aspects of Binocular Vergence Movements (1983).

Navarro, Rafael, Luis Gonzalez, and Jose L. Hernandez. “Optics of the average normal cornea from general and canonical representations of its surface topography.” JOSA A 23.2 (2006): 219–232.

Rhee, Seung Chul, Kyoung-Sik Woo, and Bongsik Kwon. “Biometric study of eyelid shape and dimensions of different races with references to beauty.” Aesthetic plastic surgery 36.5 (2012): 1236–1245.

Rohrschneider, K., et al. “Stability of fixation: results of fundus-controlled examination using the scanning laser ophthalmoscope.” German journal of ophthalmology 4.4 (1995): 197–202.

Shih, Sheng-Wen, Yu-Te Wu, and Jin Liu. “A calibration-free gaze tracking technique.” Proceedings 15th International Conference on Pattern Recognition. ICPR-2000. Vol. 4. IEEE, 2000.

Steinman, R. M., W. B. Cushman, and A. J. Martins. “The precision of gaze.” Human neurobiology 1 (1982): 97–109.

Świrski, L., & Dodgson, N. (2014, March). Rendering synthetic ground truth images for eye tracker evaluation. In Proceedings of the Symposium on Eye Tracking Research and Applications(pp. 219–222).

Tonn, Bastian6, Oliver Klaus Klaproth, and Thomas Kohnen. “Anterior surface–based keratometry compared with Scheimpflug tomography–based total corneal astigmatism.” Investigative ophthalmology & visual science 56.1 (2015): 291–298.

Wang, Jianzhong, Guangyue Zhang, and Jiadong Shi. “2D gaze estimation based on pupil-glint vector using an artificial neural network.” Applied Sciences 6.6 (2016): 174.

Yiu, Yuk-Hoi, et al. “DeepVOG: Open-source pupil segmentation and gaze estimation in neuroscience using deep learning.” Journal of neuroscience methods 324 (2019): 108307.

## References

Westheimer, Gerald. “The visual world of the new contact Lens wearer.” The Australian Journal of Optometry 46.5 (1963): 124–127.

